# Efficacy of postmenopausal estrogen replacement in SIV-infected female macaques on antiretroviral therapy

**DOI:** 10.64898/2026.09.21.746440

**Authors:** Gabriela M. Webb, Diana Takahashi, Melissa Kirigiti, Sarah R. Lindsley, Hannah Blomenkamp, Cicely Zaro, Molly Shallman, Casey McGuire, Heather Hofmeister, Cleiton Pessoa, Allyson McCullen, Matthew Humkey, Vahid Monfared, Urszula T. Iwaniec, Russell T. Turner, Lina Gao, Oleg Varlamov, Phyllis C. Tien, Jeffrey T. Jensen, Charles T. Roberts, Jonah B. Sacha, Paul Kievit, Kristin A. Sauter

## Abstract

The success of modern antiretroviral therapy (ART) has increased the life expectancy of people living with HIV to levels approaching that of people without HIV. For women living with HIV (WLWH), this means that more will survive to undergo menopause and experience the consequences of decreased ovarian hormone levels, particularly estrogen (E2). The recent change in federal guidance for use of postmenopausal hormone therapy is increasing demand for both E2-alone and E2+progestogen formulations to control adverse symptoms of menopause. The consequences and efficacy of hormone therapy in WLWH are thus an important issue for WLWH and their healthcare providers. The role of E2 replacement in postmenopausal WLWH is a significant issue because of its potential effects on control of the viral reservoir and its demonstrated beneficial metabolic effects in postmenopausal women in the general population. To address these questions, we employed a novel nonhuman primate model of postmenopausal WLWH undergoing E2 replacement. Reproductively competent female rhesus macaques were infected with simian immunodeficiency virus (SIV) and then subjected to a daily ART regimen. After complete suppression of plasma viremia, all animals underwent ovariectomy (OVX) and were then implanted with Silastic capsules containing either cholesterol vehicle or sufficient E2 to restore pre-OVX plasma levels. Plasma and cell-associated viral dynamics, immune responses, body composition, systemic and tissue-specific metabolic parameters, cytokine profiles, and parameters of bone health were followed longitudinally from baseline through 34 weeks of E2 deficiency or replacement. We found that E2 status did not significantly affect plasma or tissue viral dynamics or overall metabolic homeostasis. However, E2 replacement exerted beneficial effects on several aspects of bone health despite the chronic inflammatory state that persisted following effective ART suppression of the SIV reservoir. Our findings suggest that hormone therapy, specifically E2 replacement, offers benefit to WLWH, particularly with respect to bone loss.

## Introduction

Modern antiretroviral therapy (ART) has made HIV/AIDS a manageable chronic condition rather than a fatal disease. Initiation of ART early after infection can result in near-normal life expectancy, although 5-10 years shorter than that in persons without HIV (1, 2). Persons in whom ART has successfully suppressed viral load and restored CD4 T-cell counts exhibit an increased incidence of age-associated chronic diseases such as obesity, diabetes, and cardiovascular disease due to increased survival. However, despite effective ART suppression of viremia, people with HIV remain at greater risk for cardiometabolic disease than people without HIV due to chronic inflammation and immune dysregulation. Furthermore, current ART regimens, particularly those containing integrase inhibitors and certain nucleoside reverse transcriptase inhibitors, have been associated with metabolic pertubations (3–9).

Women living with HIV (WLWH) represent the preponderance of new cases of HIV. However, depite of significant differences in multiple aspects of HIV pathology and population demographics (10–12), including ART-associated comorbidities (13), women are still underrepresented in clinical studies (10). Effective ART regimens, in addition to increasing the risk of ART-associated metabolic disease in general, have also increased the number of WLWH who survive to undergo menopause (14). HIV itself may accelerate menopause (15–18), resulting in an additional risk for metabolic comorbidities in WLWH. There is a heightened interest in optimizing management of menopause in WLWH (19, 20), particularly since severe menopausal symptoms may reduce adherence to ART (21).

Estrogen, specifically 17-β-estradiol (E2), plays a crucial role in regulating premenopausal female physiology and metabolism (22). The relative levels of E2 in WLWH vs men living with HIV (MLWH) are thought to underlie many of the differences in HIV virology and pathology, and decreased levels of E2 during menopause (23) may contribute to the development and severity of metabolic comorbidities in WLWH. It has been previously reported that estrogen receptor-α (ERα) may function as a negative regulator of the latent reservoir (24–26), which suggests that postmenopausal E2 deficiency may also alter reservoir dynamics, specifically weakened suppression of the latent viral reservoir.

It is well established that E2 depletion immediately prior to and following menopause results in bone loss, and that hormone replacement therapy (HRT) is an effective treatment to prevent bone loss and fracture. Bone mineral density (BMD) substantially declines in perimenopause and is associated with increased fracture risk (27, 28). In addition, studies have demonstrated an adverse impact of ART itself on bone, specifically a 4% reduction in areal BMD (aBMD) following initiation of ART (29). Tenofovir disoproxil fumarate, a principal component of modern ART regimens, is known to cause bone loss in people with or without HIV (30). While newer drugs such as tenofovir alafenamide exhibit an improved bone toxicity profile, they are not benign compounds and still carry risks to bone and metabolic health (30). As a result, WLWH are subject to two sequential adverse influences on bone health; i.e., ART itself and E2 depletion with ovarian aging, which may exert additive or synergistic effects.

The nonhuman primate (NHP) model of experimental menopause induced by ovariectomy (OVX) is a well-established preclinical model for postmenopausal osteopenia. We previously demonstrated that E2 replacement exerted beneficial effects on metabolic and bone heath in this model (31). In light of the potential role of ERα in the status of the latent reservoir and the possible deleterious effects of ART per se, the current study was designed to assess the efficacy of E2 replacement on both the latent reservoir and metabolic and bone health in the context of simian immunodeficiency virus (SIV) infection and ART treatment using a novel NHP model of postmenopausal WLWH.

In addition to tissue-specific effects of SIV/ART and E2 replacement on bone, we also assessed the responses of pancreas and white adipose tissue (WAT) to SIV infection, ART, and E2 replacement based on the reported effects of E2 in these tissues. Specifically, E2 has been shown to exert multiple effects on pancreatic islet physiology and pathology and certain aspects of insulin synthesis and secretion (32–34), while WAT is also an important latent reservoir (35–37) as well as a target of E2 action (38, 39) and potentially contributes to metabolic comorbidities associated with ART through control of lipid metabolism and glucose homeostasis via secretion of multiple adipocytokines (40, 41). The role of E2 in insulin resistance (42, 43) may also be partially mediated through altered WAT function.

In the studies detailed below, we show that E2 status following experimental menopause induced by OVX does not appreciably influence ART suppression of plasma viremia or the tissue viral reservoir. In addition, we show that the beneficial effects of E2 replacement seen in our previous studies in uninfected animals are also evident in the setting of SIV and ART and residual chronic inflammation. While our studies focus on the effects of E2 per se, our findings suggest that post-menopausal HRT is appropriate for some WLWH, specifically for enhanced bone health rather than for more efficient control of the latent HIV reservoir. Our findings are particularly pertinent in the context of a recently initiated clinical trial specifically addressing the role of HRT in managing menopause in WLWH (Menopausal Hormone Therapy for Women Living with HIV (HoT); NCT06856174).

## Results

To assess the effect of E2 status on the response to SIV infection and ART, we infected 24 reproductively competent female rhesus macaques with SIVmac239M and initiated a daily ART regimen at 2 weeks post-infection (PI). At 35 weeks PI, all animals underwent ovariectomy (OVX) to achieve experimental menopause, and 3 weeks later, randomized into two groups-one (n=12) implanted with Silastic capsules containing E2 and the other (n=12) implanted with capsules containing cholesterol vehicle (hereafter termed control). E2 concentrations were monitored weekly following OVX to monitor E2 decline and assess implant performance. As described below, E2 implants effectively maintained replacement levels in the desired physiological range. As also described below, one animal was excluded from the study due to detectable estrogen levels post-OVX. A schematic of the study design is shown in **Figure 1A**.

**Figure 1.**
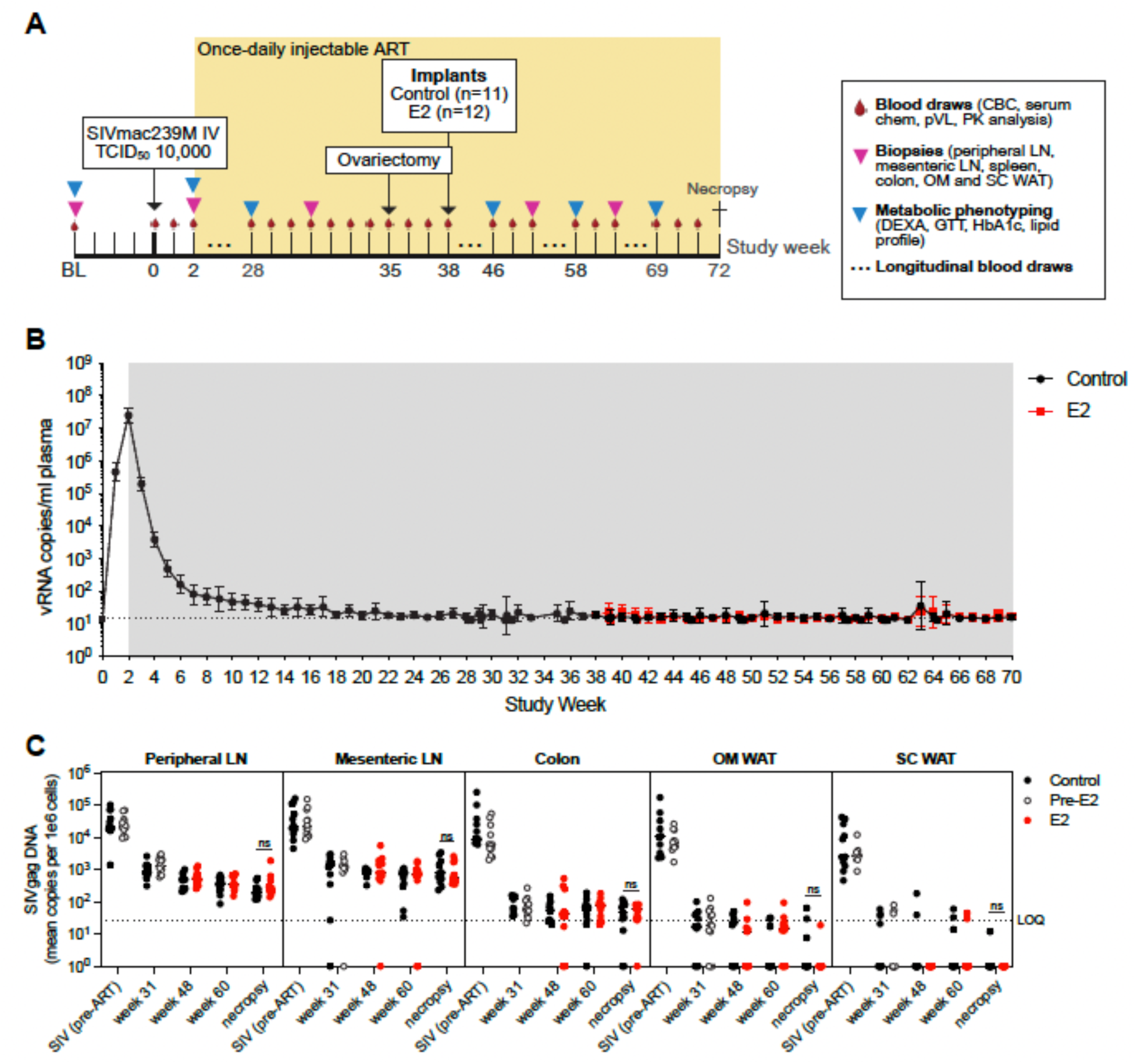
Experimental timeline and plasma and cell-associated viral DNA dynamics during SIV infection, ART suppression, and E2 replacement. **A.** Schedule of experimental procedures and assessments. **B.** SIV RNA copies per ml of plasma over the study time course. Limit of quantitation (LOQ) is denoted by the horizontal dotted line. Black and red points starting at week 37 PI represent control and E2-replaced groups, respectively. There were no significant differences between the average plasma viral loads in the groups of animals destined to receive vehicle or E2-containing implants (data not shown). **C.** Cell-associated SIV DNA in peripheral lymph node, colon, and the SVF of OM and SC WAT over the study time course. As in panel B, black and red points starting at week 37 PI represent control and E2-replaced groups, respectively, and there were no significant differences between the average cell-associated viral loads in the groups of animals destined to receive vehicle or E2-containing implants. All data are means ± SEM.

### Effect of E2 replacement on viral dynamics

Longitudinal changes in plasma viral loads are shown in **Figure 1B**. Plasma SIV RNA levels increased rapidly until daily ART was instituted at 2 weeks PI and then declined until full suppression was achieved around 24 weeks PI. Following OVX and implant placement at 35 and 38 weeks PI, respectively, no differences in plasma viral loads were seen between the control and the E2-replaced replaced groups. Thus, E2 status did not influence plasma viral dynamics. Cell-associated SIV DNA levels, representing the viral reservoir, are shown in **Figure 1C**. Similar to what was observed with plasma viral loads, cell-associated SIV DNA levels in peripheral and mesenteric lymph nodes, colon, and the stromovascular fraction (SVF) of omental (OM) and subcutaneous (SC) WAT were significantly reduced by 31 weeks PI, and remained at reduced levels in both control and E2-replaced groups throughout the remainder of the experimental time course. These data suggest that E2 status does not affect steady-state plasma viral load or tissue cell-associated SIV DNA levels following ART treatment.

### Effect of E2 replacement on body composition

**Table 1** shows the characteristics of the animals at baseline (week −4) and immediately prior to OVX at week 35 PI. As described below, one animal that retained detectable E2 levels following OVX was removed from the study, leaving 11 animals that received cholesterol vehicle implants and 12 that received E2 implants. As shown in the left half of the table, there were significant decreases in HBA1c, total cholesterol, and LDL and HDL levels after SIV infection and ART at 34 weeks PI. However, as shown in the right half of the table, there were no differences in the animals randomized to receive cholesterol vehicle vs E2 implants. **Figure 2** shows the % change in body weight (BW) and total, lean, and fat mass determined by dual-energy X-ray (DXA) scanning over the entire experimental time course. There were no significant changes in any of these parameters other than a transient decrease in lean mass at 2 weeks PI compared to baseline (**Figure 2C**) and decreased fat mass in E2-replaced animals at 58 and 70 weeks PI compared to control (**Figure 2D**).

**Figure 2.**
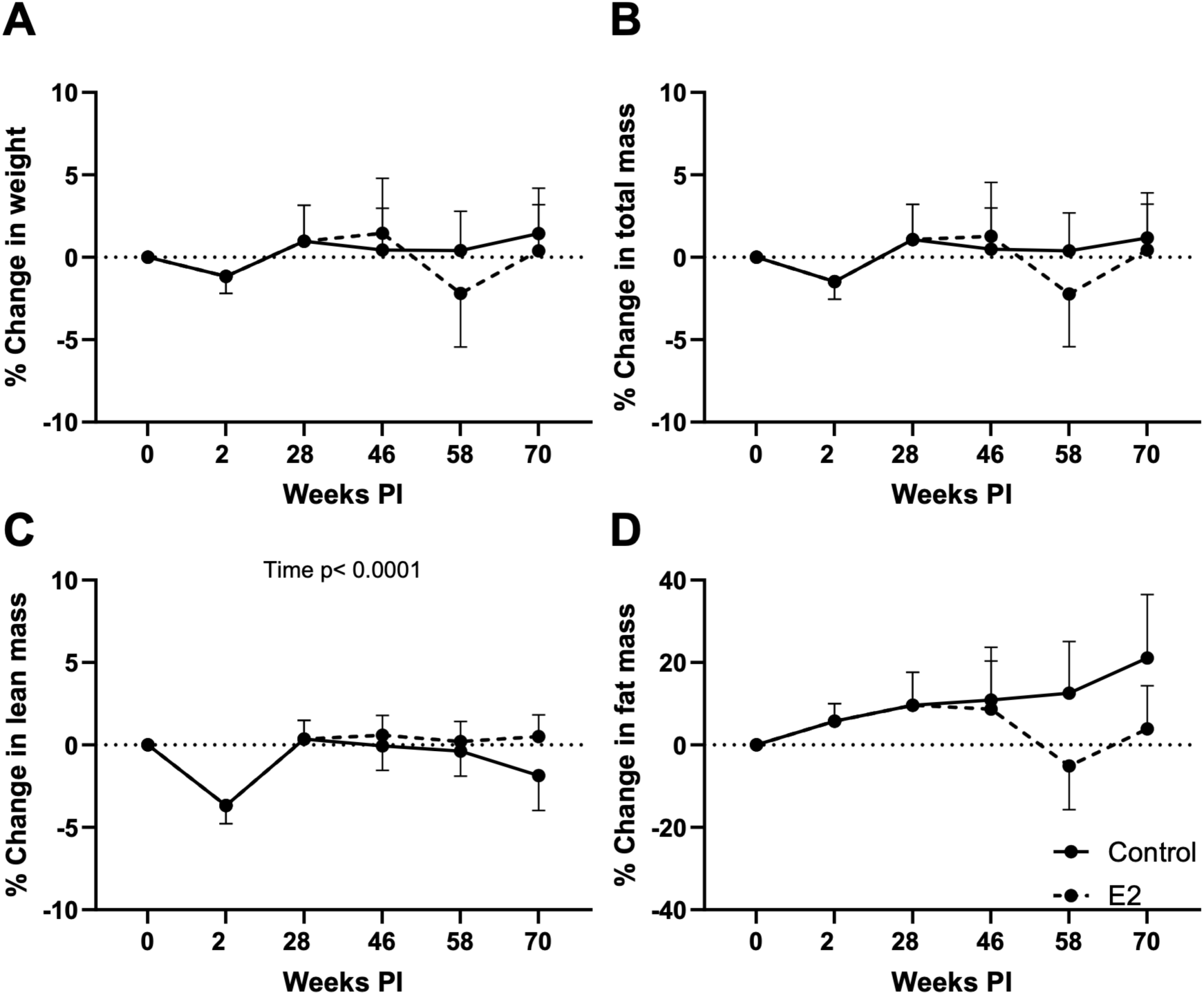
Percent change in (A) body weight (BW) and (B) total, (C) lean, and (D) fat mass during SIV infection, ART suppression, and E2 replacement. B. % change in body weight (BW) over the study time course. All data are means ± SEM. Significance determined by mixed-effects analysis with post-hoc pair-wise comparisons.

**Table 1.**
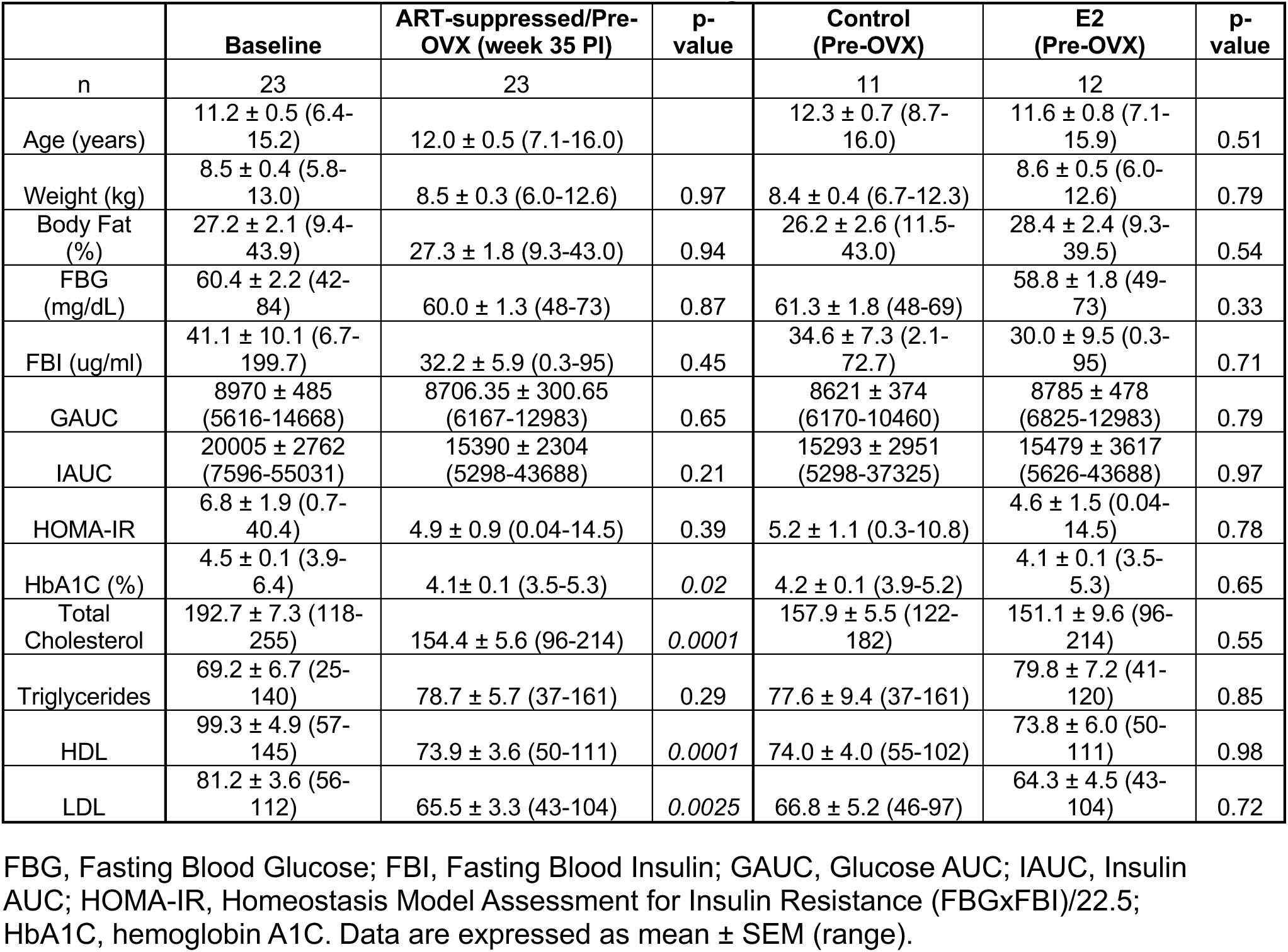
Baseline characteristics of the experimental groups.

### E2 replacement does not modulate T cell and humoral immune responses

SIV-specific T cell and humoral responses were assessed by ELISpot and env-specific binding antibody assays, respectively. As shown in **Figure 3A**, we assessed the magnitude and breadth of SIV-specific T-cell responses in blood following OVX and implant placement via ELISpot and, although there was an increased cumulative ELISpot response at weeks 68 and at necropsy in the E2 group, these differences were not statistically significant. As shown in **Figure 3B**, we also assessed the levels of anti-SIVmac239 gp140-binding antibodies over the entire experimental time course, which increased rapidly following infection, peaking at 4 weeks post-infection (2 weeks post-ART), and remained elevated for the remainder of the study period. There was no significant difference in anti-SIVmac239 gp140-binding antibodies between the control and E2-replaced groups.

**Figure 3.**
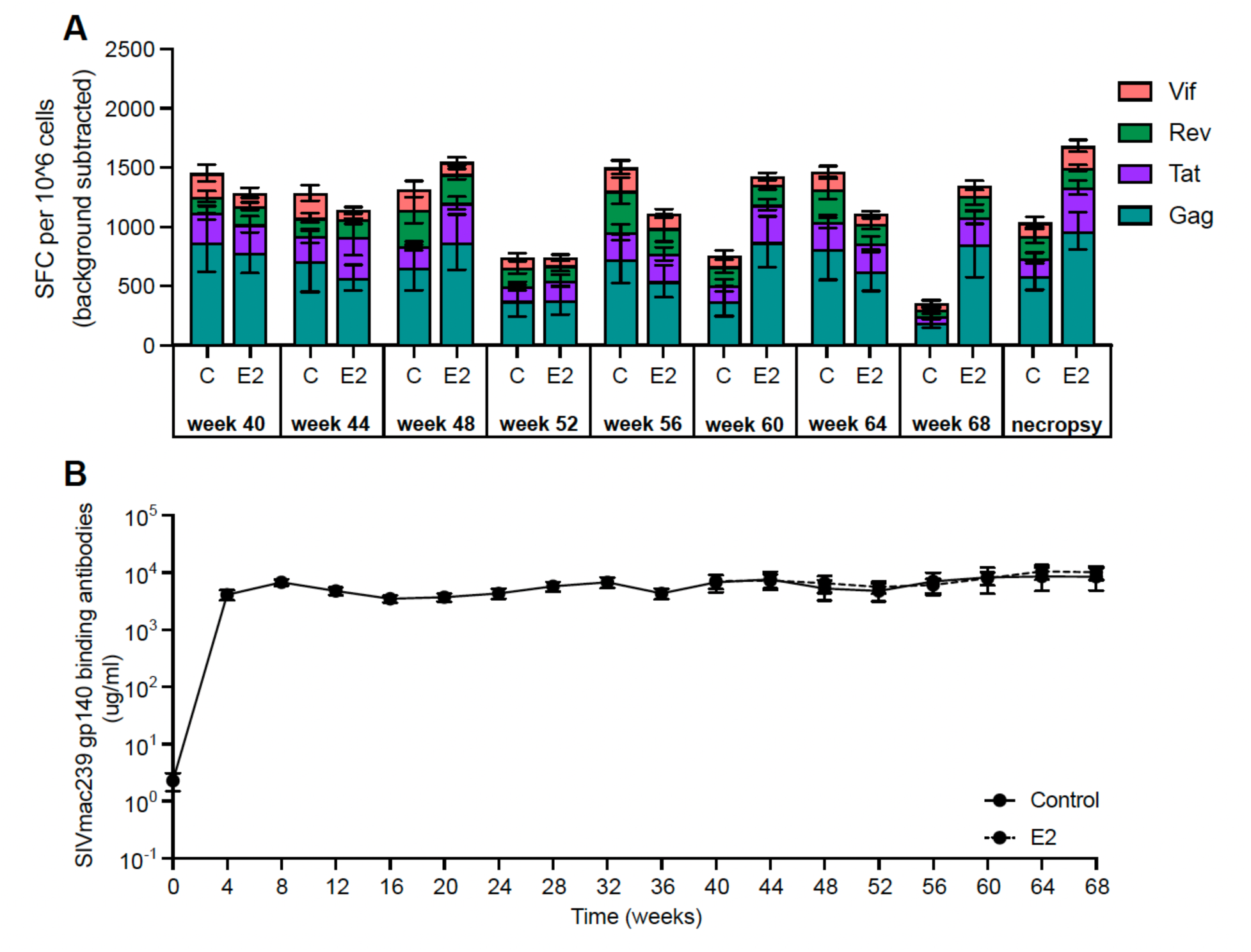
ENV-binding antibody responses and SIV-specific T-cell responses during SIV infection, ART suppression, and E2 replacement. A. Longitudinal change in SIVmac239 gp140-binding antibodies in plasma following OXV and implant placement. B. IFNγ spot-forming cells (SFC) per 10^6^ mononuclear cells were quantified by ELIspot upon stimulation with subpools of peptides spanning Gag, Tat, Rev, and Vif. All data are means ± SEM.

### E2 replacement does not significantly influence circulating and WAT immune cell profiles

**Figure 4** shows the proportions of CD4+ and CD8+ T cells in whole blood (WB; **Figure 4A**) and the SVF of OM (**Figure 4B**) and SC WAT (**Figure 4C**) over the experimental time course. SIV infection, ART treatment, and subsequent E2 status did not alter these parameters in any of these compartments, other than a transient increase in CD8+ T cells at 2 weeks PI in whole blood (lower panel in **Figure 4A**). These results in WAT differ from those we previously described in SIV-infected, ART-treated males, in which there was a significant decrease in CD4+ T cells and a significant increase in CD8+ T cells in both OM and SC WAT after SIV infection that was not resolved by ART (44). We also assessed T cell activation as assessed by Ki67, CD38, and CD69 staining of CD4+ and CD8+ T cells in WB (**Figure 5A**), OM WAT (**Figure 5B**), and SC WAT (**Figure 5C**), with the only changes being transient increases in Ki67+ and CD38+CD8+ T cells in OM and SC WAT during initial SIV infection (2 weeks PI) that were reversed following ART.

**Figure 4.**
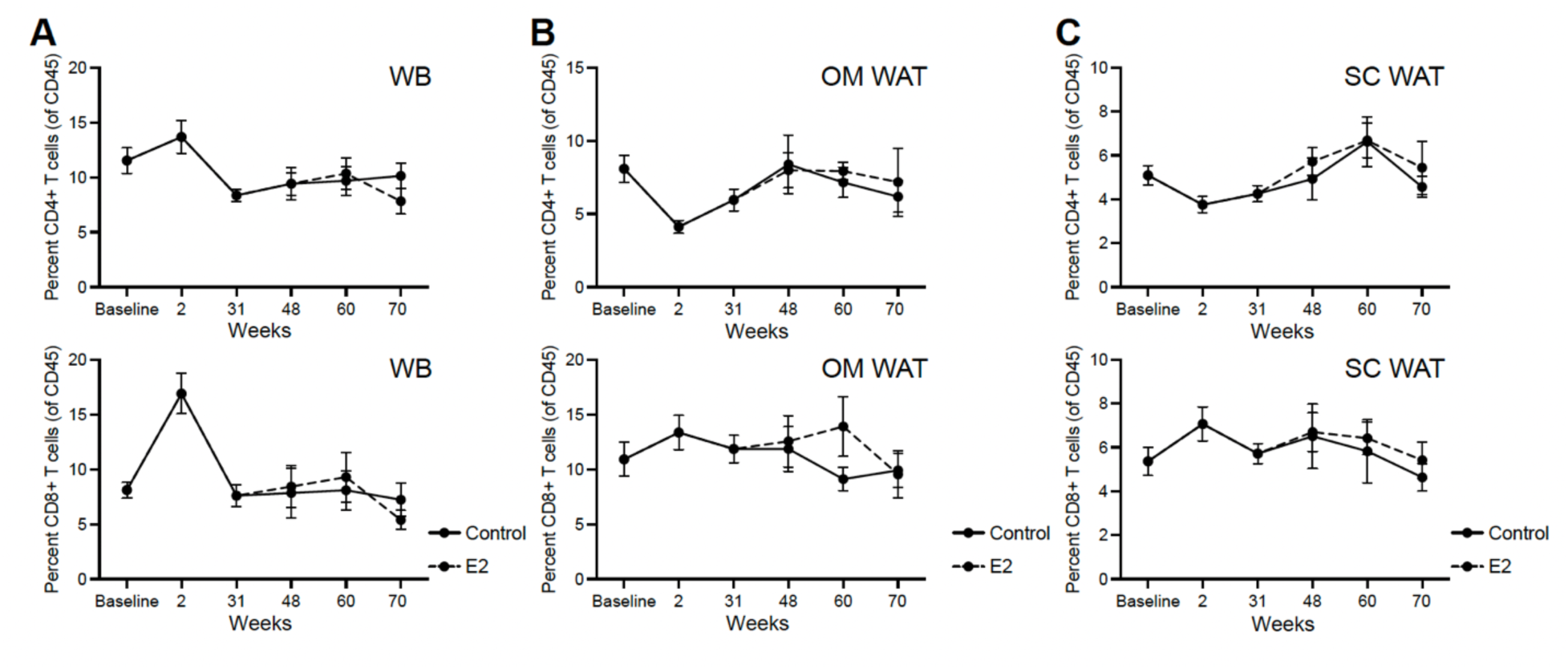
Effect of SIV infection, ART suppression, and E2 replacement on CD4+ and CD8+ T-cell frequencies in (A) whole blood (WB) and (B) OM and (C) SC WAT. WB monocytes and the stromovascular fraction of OM and SC WAT were analyzed by flow cytometry using the gating strategy and antibodies described in Figure 16 and Table 4. All data are means ± SEM.

**Figure 5.**
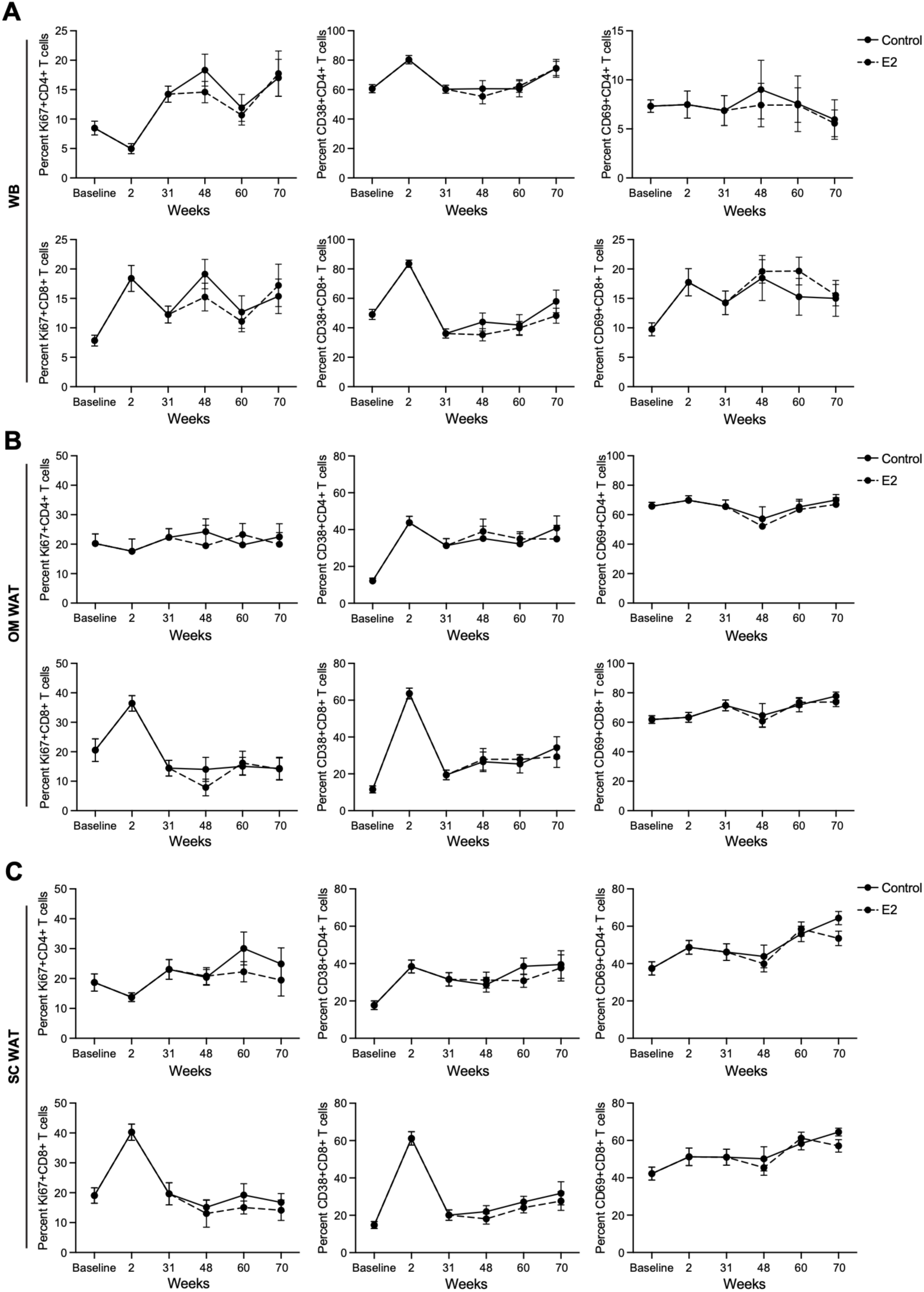
Effect of SIV infection, ART suppression, and E2 replacement on CD4+ and CD8+ T-cell activation in (A) WB and (B) OM and (C) SC WAT. Proportions of Ki67, CD36, and CD69-positive CD4+ and CD8+ T-cells in WB monocytes and the stromovascular fraction of OM and SC WAT were analyzed by flow cytometry using the gating strategy and antibodies described in Figure 16 and Table 4. All data are means ± SEM.

### Effect of E2 replacement on parameters of systemic metabolic control

Intravenous (iv) glucose tolerance tests (ivGTTs) were used to assess the effect of E2 replacement on in vivo insulin secretion and glucose clearance. As shown in **Figure 6A**, glucose excursion dynamics during an ivGTT were similar between baseline and immediately prior to necropsy in both the E2-replaced and control groups, although the percent change in glucose area under the curve (GAUC) significantly increased in both groups following ART initiation, with values slightly higher in the E2-deficient group (**Figure 6B**), suggesting that E2 replacement tended to maintain insulin sensitivity. Somewhat unexpectedly, insulin secretion during an ivGTT was decreased in both experimental groups at 70 weeks PI compared to baseline, although more pronounced in the E2-replaced group (**Figure 6C**); this effect was also evident in a significant decrease in the percent change in insulin AUC (IAUC), which was again more evident in the E2-replaced group (**Figure 6D**).

**Figure 6.**
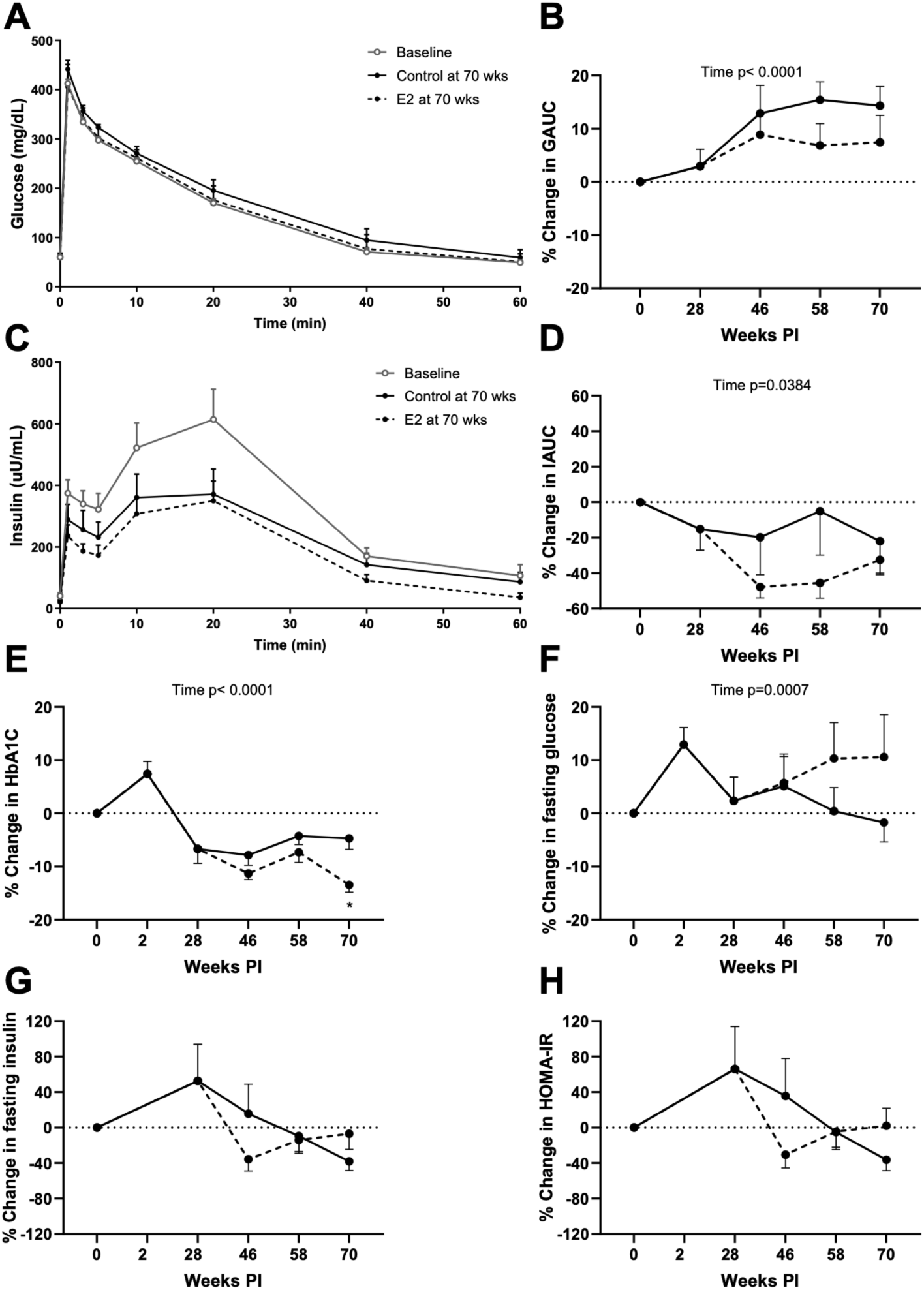
Effect of SIV infection, ART suppression, and E2 replacement on glucose metabolism. A. Glucose excursions in ivGTTs performed at baseline and in the control and E2-replaced groups at 70 weeks PI. B. Glucose AUC change (%) in longitudinal ivGTTs in the control and E2-replaced groups. C. Insulin excursions in ivGTTs performed at baseline and in the control and E2-replaced groups at 70 weeks PI. D. Insulin AUC change (%) in longitudinal ivGTTs in the control and E2-replaced groups. E. Longitudinal changes (%) in HbA1c in the control and E2-replaced groups. F. Longitudinal changes (%) in fasting glucose in the control and E2-replaced groups. G. Longitudinal changes (%) in fasting insulin in the control and E2-replaced groups. H. Longitudinal changes (%) in HOMA-IR in the control and E2-replaced groups. Data are means ± SEM. Significance determined by mixed effects analysis with post-hoc pair-wise comparisons. *, p < 0.05

Longitudinal changes were also seen in HbA1C and fasting blood glucose, with a significant decrease in HbA1c in the E2-replaced group at 70 weeks PI (**Figure 6E,F**), while no significant longitudinal changes were seen in fasting insulin levels or Homeostatic Model Assessment of Insulin Resistance (HOMA-IR) values (**Figure 6G,H**).

### Effect Of E2 replacement on β-cell function

To more directly assess the potential effects of E2 status on β-cell function, we evaluated glucose-stimulated insulin secretion (GSIS) and respiratory capacity in isolated pancreatic islets obtained at necropsy from a subset of E2-replaced and deficient animals (n=6 control and 6 E2-replaced). As shown in **Figure 7**, there were no differences in first-or second-phase insulin secretion (**Figure 7A**) or IAUC (**Figure 7B**) between the control and E2-replaced groups. Islet respiratory assessments are shown in **Figure 8**, which showed that islets from E2-replaced animals exhibited significantly reduced maximal and spare oxygen consumption compared to control, indicating impaired mitochondrial energy generation.

**Figure 7.**
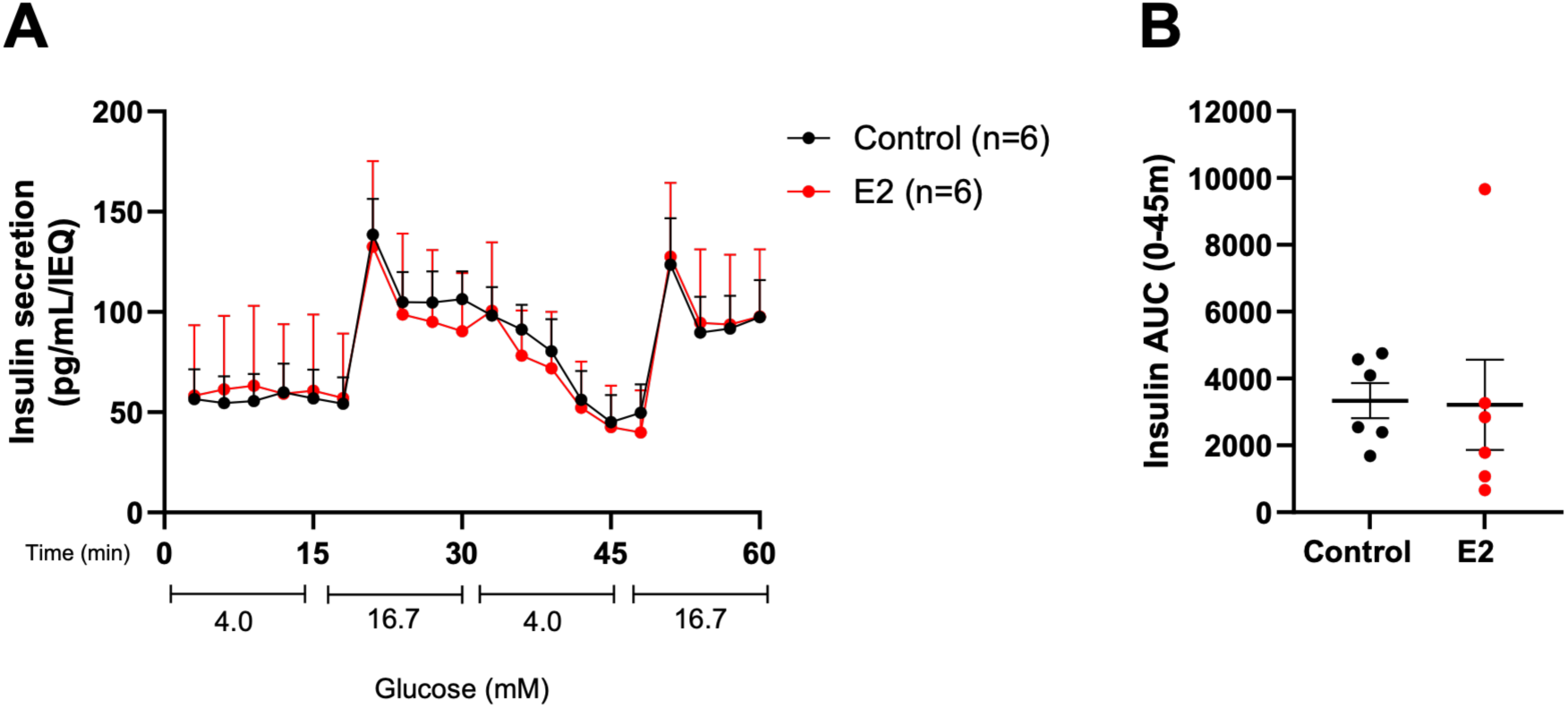
Effect of SIV infection, ART suppression, and E2 replacement on glucose-stimulated insulin secretion GSIS). A. Perifusion analysis of GSIS in islets isolated from control and E2-replaced groups. B. Insulin AUC values in a single high-glucose stimulation in individual islet isolations from control and E2-replaced animals. Data are means ± SEM.

**Figure 8.**
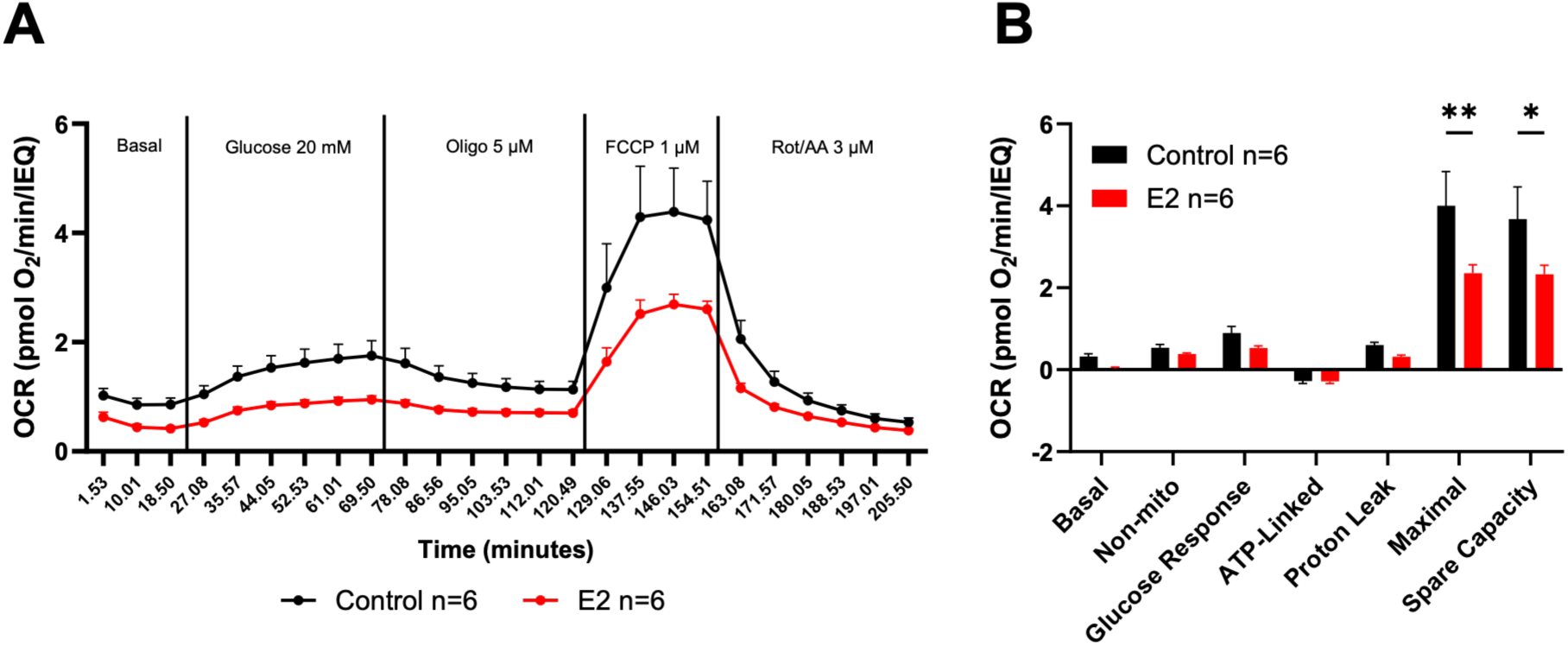
Effect of SIV infection, ART suppression, and E2 replacement on islet oxygen consumption. A. Oxygen consumption rate (OCR) in islets isolated from control and E2-replaced groups during treatment with glucose, oligomycin, FCCP, and Rot/AA. B. Differences in indicated OCR parameters in control and E2-replaced islets derived from the data of panel A. All data are means ± SEM. Significance was determined by 2-way ANOVA with multiple comparisons *, p<0.05; **, p<0.01.

### Effect of E2 replacement on plasma lipids

We also assessed plasma lipid levels over the experimental time course. As shown in **Figure 9A-D**, longitudinal changes were seen in plasma triglycerides, total cholesterol, HDL, and LDL, with cholesterol and HDL levels dropping significantly after ART initiation and remaining lower in both E2-replaced and deficient groups (**Figure 9B,C**). LDL levels also decreased following infection and ART initiation but trended back toward baseline levels in the control group while remaining depressed in the E2 group (**Figure 9D**).

**Figure 9.**
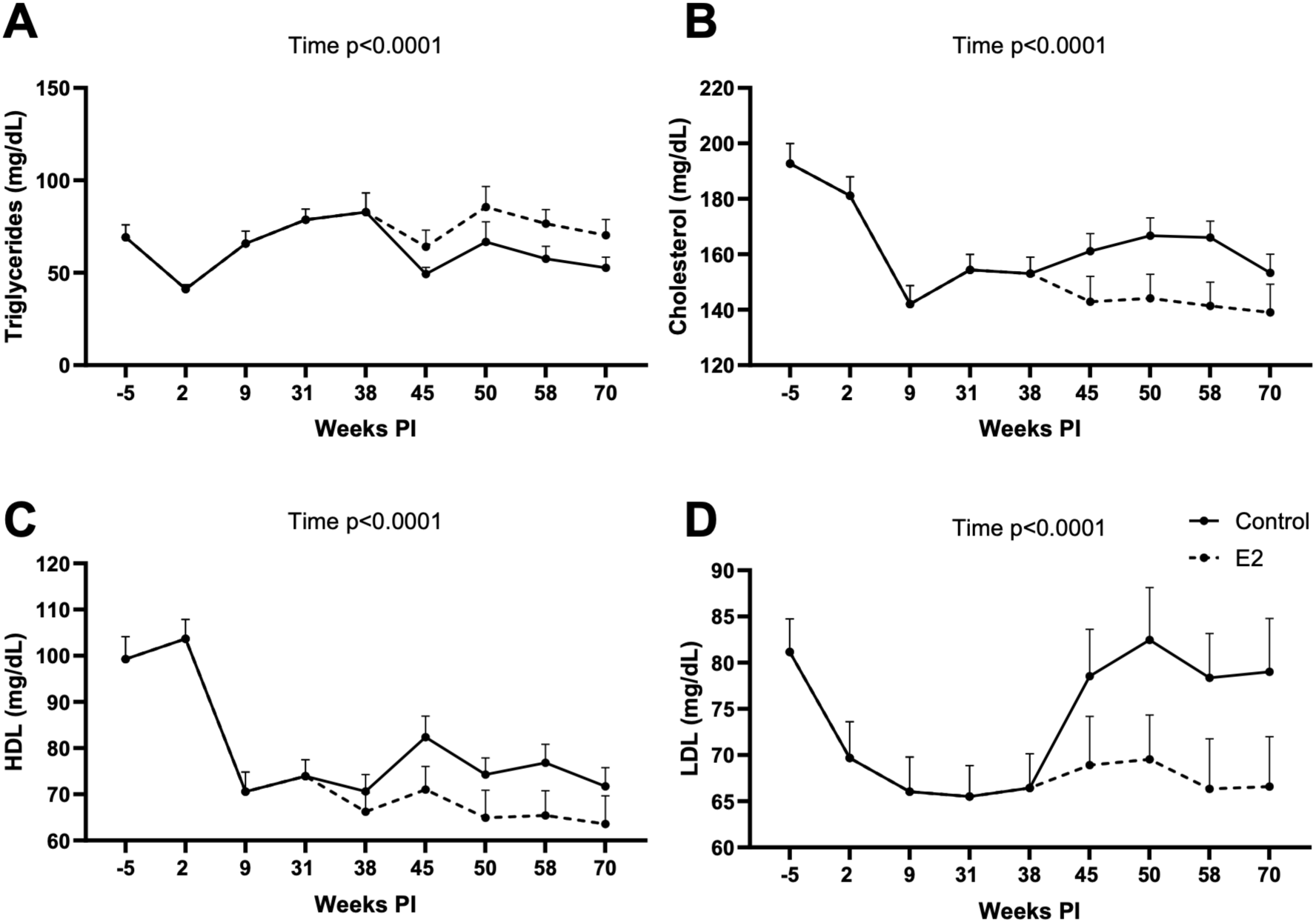
Effect of SIV infection, ART suppression, and E2 replacement on plasma lipid levels. Longitudinal changes in (A) triglycerides, (B), cholesterol, (c) HDL, and (D) LDL in the control and E2-replaced groups. All data are means ± SEM. Significance determined by mixed effects analysis with post-hoc pair-wise comparisons.

### Effect of E2 replacement on WAT morphology and function

In view of the known effects of E2 on WAT, we investigated the effect of E2 deficiency and replacement on adipocyte size (area), WAT extracellular matrix (ECM) thickness (a surrogate measure of WAT expandability) and plasma levels of the major adipokines adiponectin and leptin. As shown in **Figure 10 A,B**, there was a significant increase in adipocyte area in both OM and SC WAT over the experimental time course, but this was not different between the control and E2-replaced groups. There was a significant but transient increase in ECM thickness at peak infection (2 weeks PI) but ECM thickness was unchanged thereafter in both control and E2-replaced groups (**Figure 10C,D**). As shown in **Figure 10E-G**, there were significant longitudinal decreases in the circulating levels of adiponectin and leptin and the adiponectin:leptin ratio (ALR; a biomarker of cardiometabolic risk) following SIV infection and ART, but there was no additional effect of E2 status.

**Figure 10.**
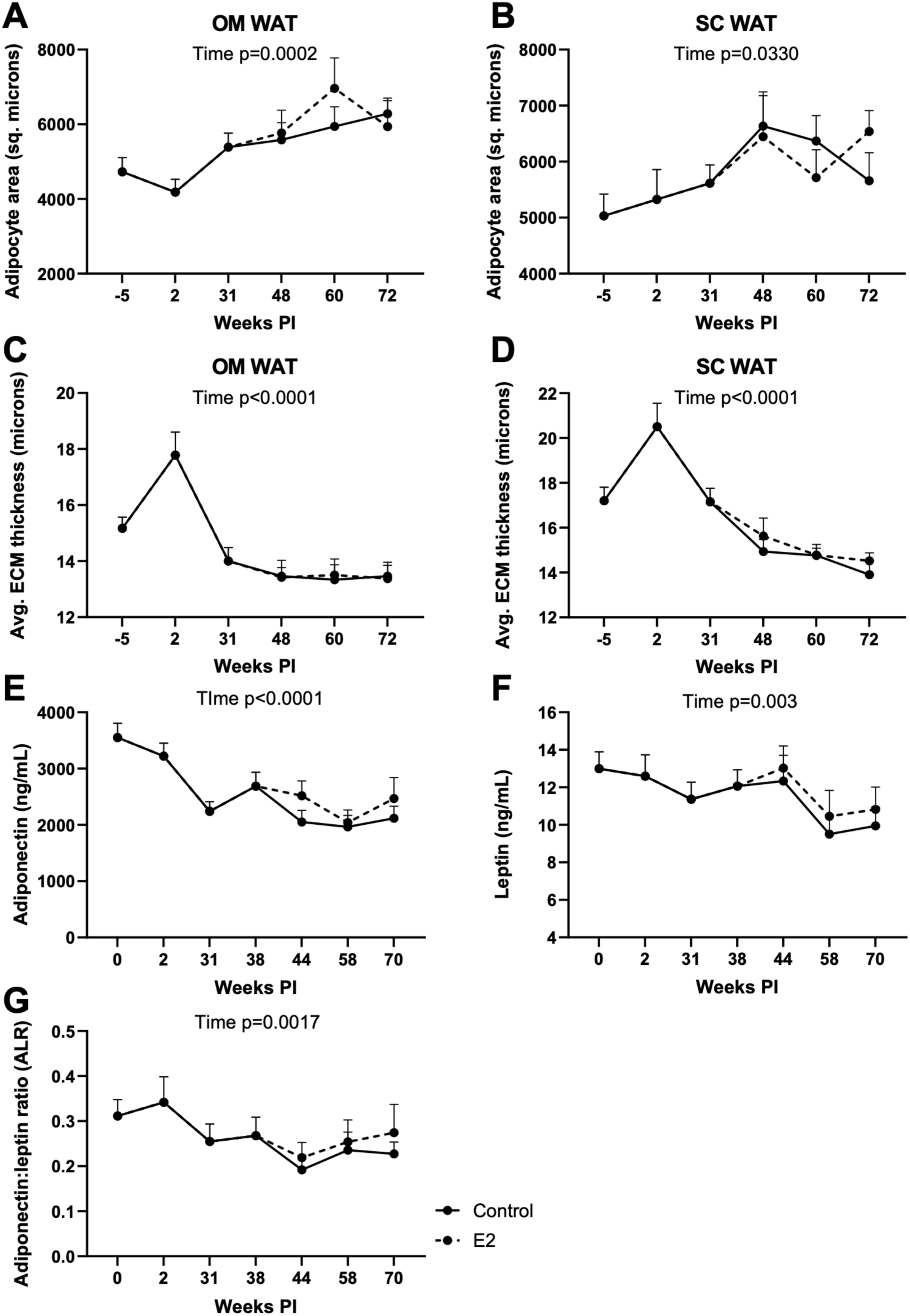
Effect of SIV infection, ART suppression, and E2 replacement on adipocyte size, pericellular extracellular matrix thickness, and adipocytokine levels. Average adipocyte area in (A) OM and (B) SC WAT and average inter-adipocyte extracellular matrix (ECM) thickness in (C) OM and (D) SC WAT over the experimental time course. Longitudinal changes in (E) adiponectin, (F), leptin, and (G) adiponectin:leptin ratio (ALR) in the control and E2-replaced groups. All data are means ± SEM. Significance determined by mixed effects analysis with post-hoc pair-wise comparisons.

Overall, these findings suggest that SIV infection and ART did not have a major impact on metabolic homeostasis over the experimental time course but did result in potential increased cardiometabolic risk as reflected by the decrease in adiponectin levels and the ALR, similar to what we have recently reported in SIV-infected, ART-treated male macaques (44). However, these effects were largely unaffected by E2 status.

### SIV-induced increases in plasma cytokines are not prevented by ART or E2 replacement

A subset of plasma samples from E2-replaced animals (n=6) were analyzed using the NULISA cytokine profiling platform (NULISAseq inflammation panel 250, Alamar Biosciences, Fremont, CA). The principal component analysis (PCA) plot in **Figure 11A** shows that the most variation in target cytokine expression occurred at peak infection at 2 weeks PI (green area) compared to baseline (red area). The expression patterns at subsequent timepoints corresponding to full suppression (31 weeks PI; blue area), three weeks post-OVX (38 weeks PI; orange area), and following 20 weeks of E2 replacement (58 weeks PI; purple area), were overlapping but still distinct from the patterns at baseline and peak viremia, suggesting that SIV infection resulted in a significant change in the plasma cytokine profile that did not return to the baseline profile following subsequent ART and E2 replacement. A significance threshold plot of analytes exhibiting increasing degrees of change over the experimental time course is shown in **Figure 11B**. The majority of these exhibited the highest abundance at peak viremia at 2 weeks PI, followed by a variable decrease following ART initiation. However, numerous cytokiness remained at above-baseline levels following ART initiation, OVX, and E2 replacement as shown in the volcano plot in **Figure 11C**, suggesting that SIV infection results in a chronic inflammatory state in spite of effective suppression of plasma viremia.

**Figure 11.**
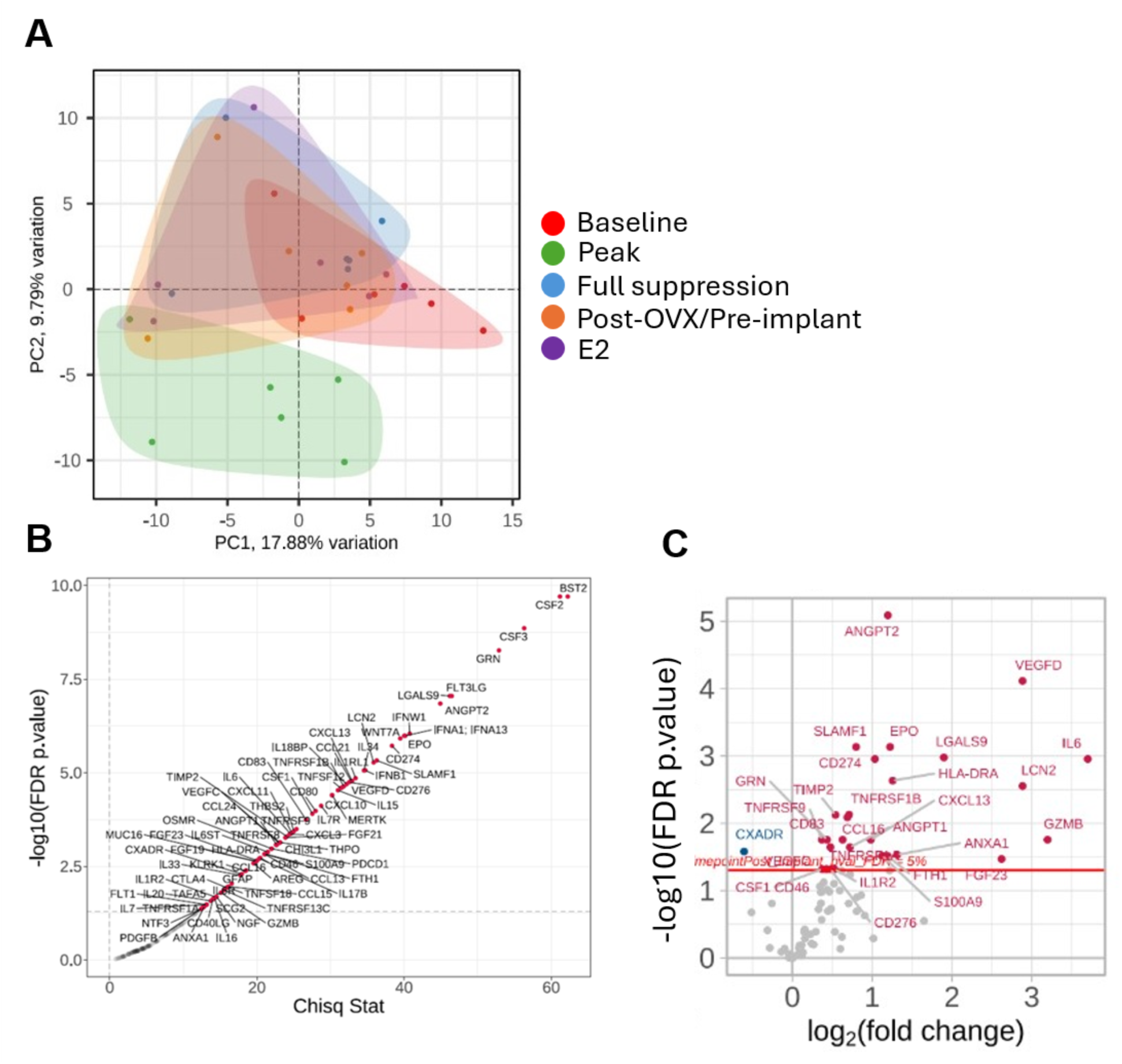
Effect of SIV infection, ART suppression, and E2 replacement on plasma cytokine profiles. A. Principal component analysis (PCA) of temporally regulated cytokines analyzed by NULISA. B. Significance threshold plot of cytokines exhibiting differential abundance over the experimental time course. C. Volcano plot of cytokines exhibiting differences in abundance between baseline and final timepoint values. Red datapoints represent analytes whose levels at week 58 PI were significantly greater that at baseline while blue datapoint is single analyte whose levels at week 58 PI were significantly lower than at baseline.

There were three general patterns of change in plasma cytokine levels illustrated in **Figure 12**. The majority of factors that exhibited the greatest degree of change over the experimental time course displayed the pattern represented by BST2, CSF2, CSF3, CXCL10 and CXCL11, in which there was a maximal level at peak viremia at 2 weeks PI followed by a return to baseline levels at week 31 PI after 29 weeks of ART (**Figure 12A**). The second pattern was represented by LGALS9 and PD-L1/CD274, in which there was a maximum at peak viremia at 2 weeks PI that decreased by 31 weeks PI following 29 weeks of ART but not to baseline levels (**Figure 12B**), representing persistent inflammation in spite of effective suppression of plasma viremia. The third pattern, represented by factors including LCN2, FGF-23, IL6, and VEGF-D (**Figure 12C**), was an increase following ART initiation that persisted after OVX and 20 weeks of E2 replacement (LCN2 and FGF-23) or a further increase following OVX (IL6 and VEGF-D). The patterns seen in **Figure 12B and 12C** define a persistent inflammatory state that is independent of effective ART.

**Figure 12.**
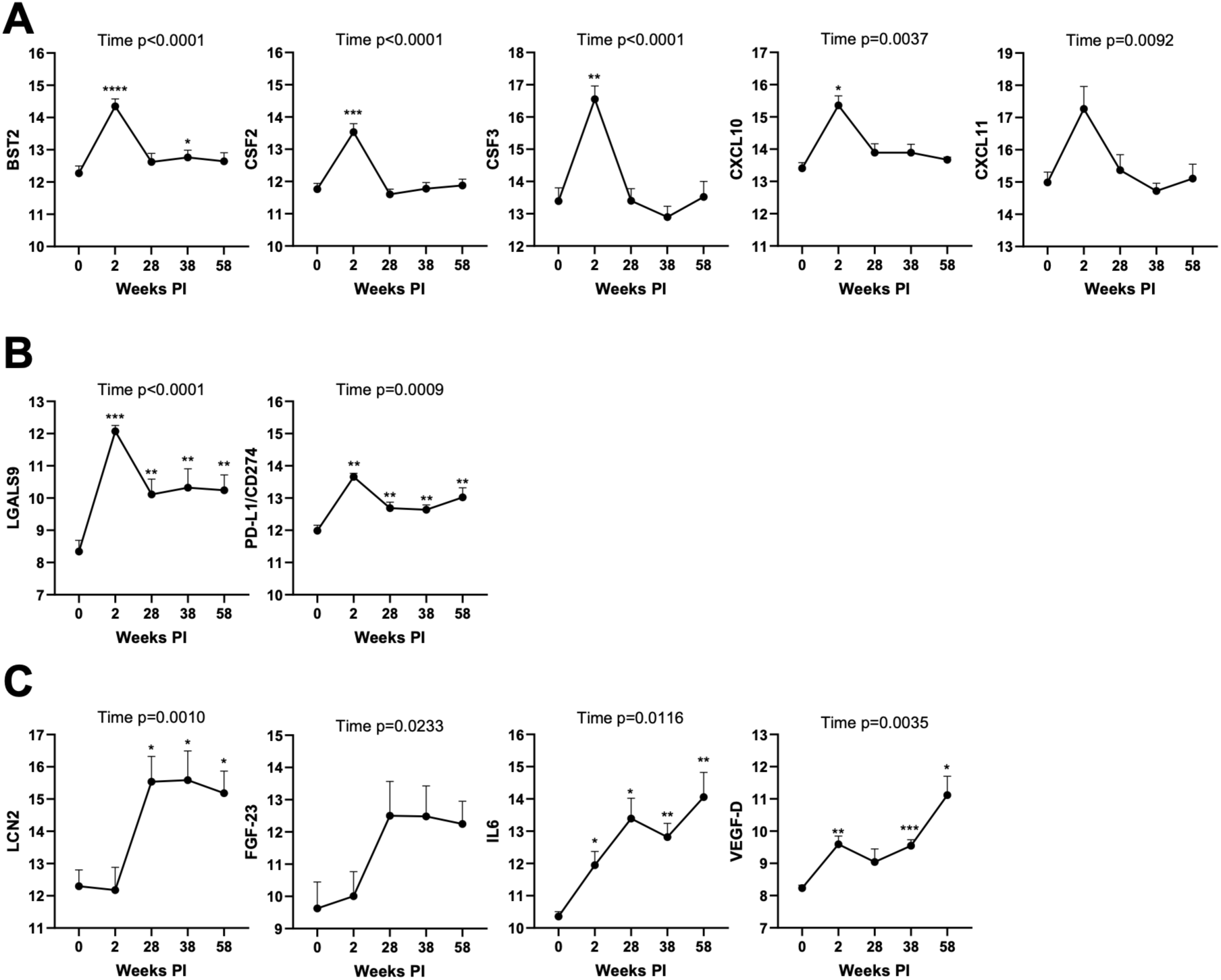
Categories of longitudinal expression patterns of selected cytokines during SIV infection, ART, OVX, and E2 replacement. A. Cytokines exhibiting a maximum at peak viremia followed by return to baseline levels. B. Cytokines exhibiting a maximum at peak viremia followed by a decrease but not to baseline levels. C. Cytokines exhibiting an increase following ART initiation with a sustained higher level (LCN2 and FGF-23) or a further increase following OVX and E2 replacement (IL6 and VEGF-D). All data are means ± SEM. Significance determined by one-way ANOVA analysis with post-hoc multiple comparisons. *, p < 0.05; **, p < 0.01; ****, p < 0.0001.

### Effects of E2 replacement on bone

Whole-body bone health can be inferred from parameters such as bone mineral content (BMC) and bone mineral density (BMD) determined by DXA. Whole-body BMC and BMD values (as percent change from baseline) remained relatively constant over most of the study period. However, at 70 weeks PI, the percent change in whole-body BMC and BMD values in the E2 replacement group was significantly higher than in the control group (**Figure 13A,B**). Specifically, there was a 2.04±0.56% increase in BMC in the E2 group vs a −0.24±1.16% decrease from baseline in BMC in the control group, while there was a 5.52±1.12% increase in BMD from baseline in the E2 group vs a 1.44±1.26% increase in BMD from baseline in the control group.

**Figure 13.**
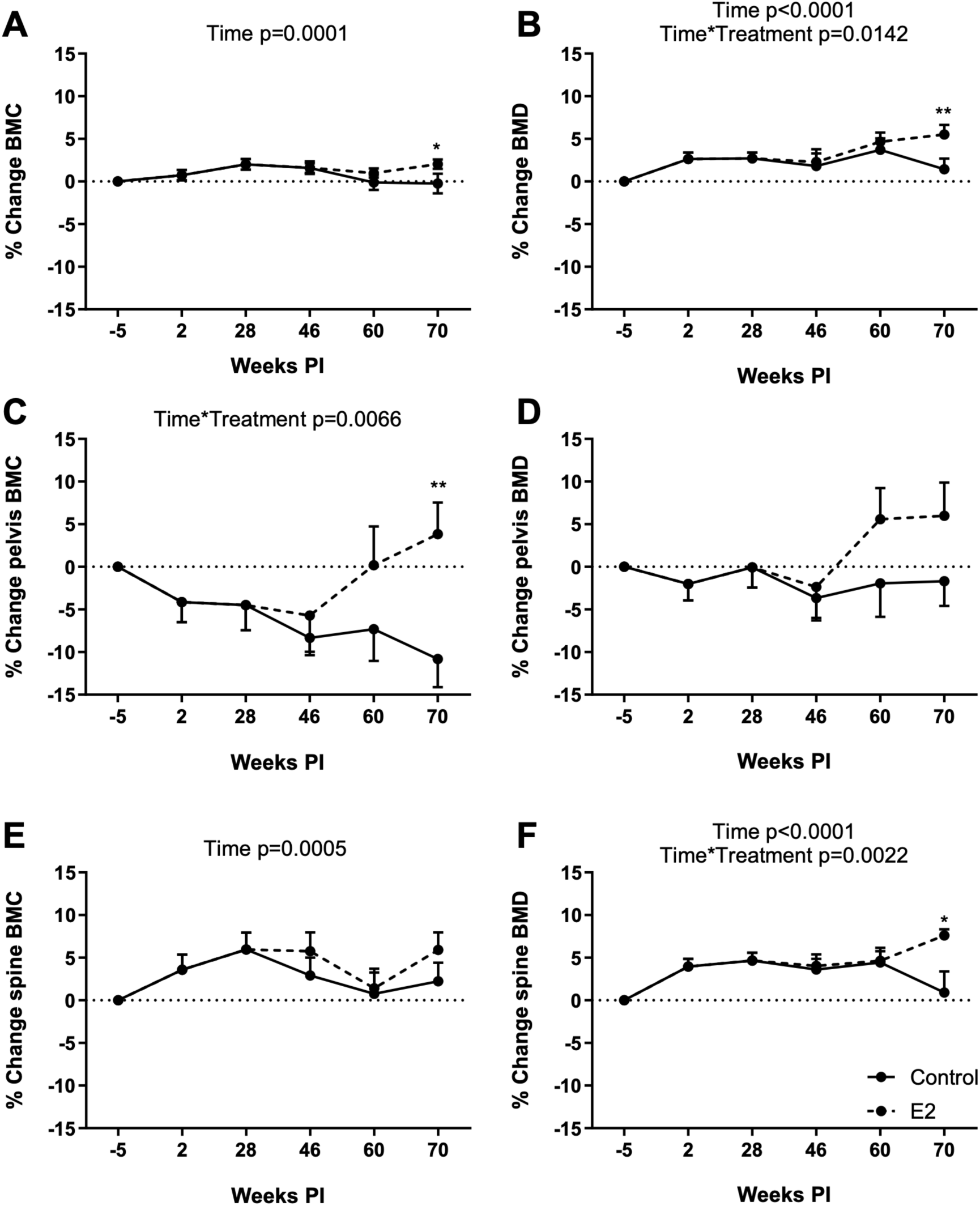
Effect of SIV infection, ART suppression, and E2 replacement on bone parameters. Longitudinal changes (%) in the control and E2-replaced groups for (A) overall Whole, (B) overall BMD, (C) pelvis BMC, (D) pelvis BMD, (E) spine BMC, and (F) spine BMD measured using DXA. All data are means ± SEM. Significance determined by mixed effects analysis with post-hoc pair-wise comparisons. *, p < 0.05; **, p < 0.01

To determine which regions contributed to these changes in whole-body BMC and BMD, we analyzed those specific areas of the skeleton typically examined in post-menopausal women. There was a progressive loss in pelvis BMC that was significant in the control group at 70 weeks PI versus baseline (−10.80±3.32%) and this loss of pelvic BMC was prevented by E2 replacement (3.83±3.71% increase from baseline; **Figure 13C**). Pelvis BMD did not change over the time course in the control group but trended above baseline in the E2-replaced group following E2 replacement (**Figure 13D**). Changes in spine BMC were not different between the two groups (**Figure 13E**), while changes in spine BMD in the E2 replacement group were significantly elevated compared to control at 70 weeks PI (7.61±0.73% increase from baseline in the E2 group vs a 0.92±2.47% increase from baseline in the control group; **Figure 13 F**).

To evaluate bone architecture, microcomputed tomography (μCT) analysis of the distal femur (cancellous and cortical bone), proximal tibia (cancellous bone), and lumbar vertebra (cancellous bone) was performed at necropsy at 72 weeks PI. E2-replaced females had significantly higher bone volume/tissue volume (BV/TV; 13.9±1.1 vs 10.7±0.8 %), connectivity density (4.9±0.5 vs 3.4±0.4 mm^-3^), and trabecular number (1.5±0.06 vs 1.3±0.06 mm^-1^) and significantly lower trabecular spacing (694.6±32.3 vs 795.0±34.8 µm) with no change in trabecular thickness (144.1±5.5 vs 142.6±5.8 µm) in distal femur metaphysis (**Figure 14A-E**). The differences in cancellous bone in the distal femur in response to treatment can be readily appreciated in representative μCT images (**Figure 15**). We also assessed cortical bone in the femur. Unlike the effects of E2 on cancellous bone, cortical bone parameters (cross-sectional volume, cortical volume, marrow volume, cortical thickness, and polar moment of inertia) were unaffected (**Table 2**). In addition, significant differences in cancellous bone in response to treatment were not detected in proximal tibia metaphysis (**Table 3**).

**Figure 14.**
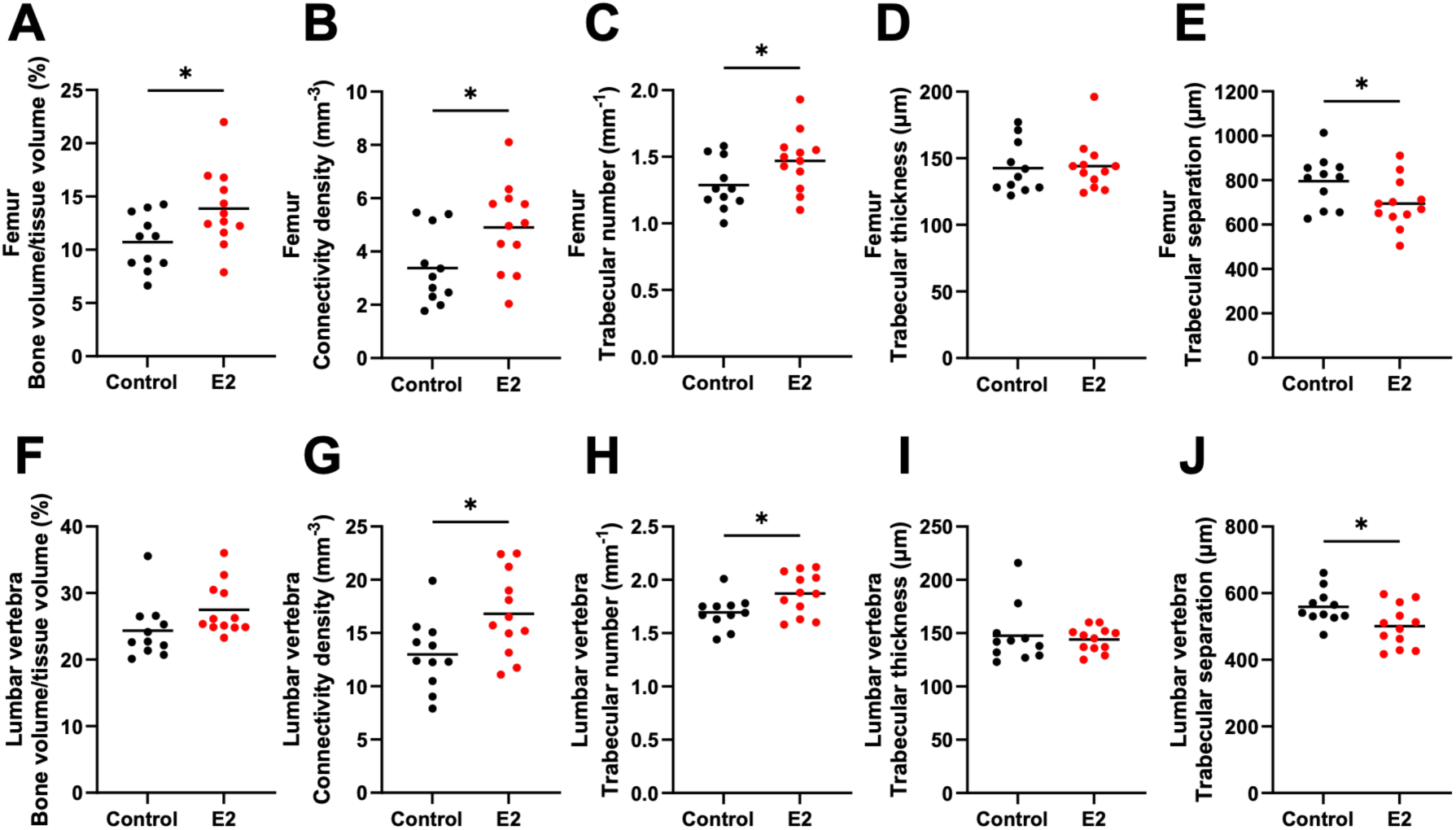
Effect of SIV infection, ART suppression, and E2 replacement on femoral and lumbar vertebra architecture. Cancellous bone of interest evaluated via microcomputed tomography in the metaphysis of distal femur and lumbar vertebra. All data are means ± SEM. Significance determined by Welch’s t test. *, p<0.05.

**Figure 15.**
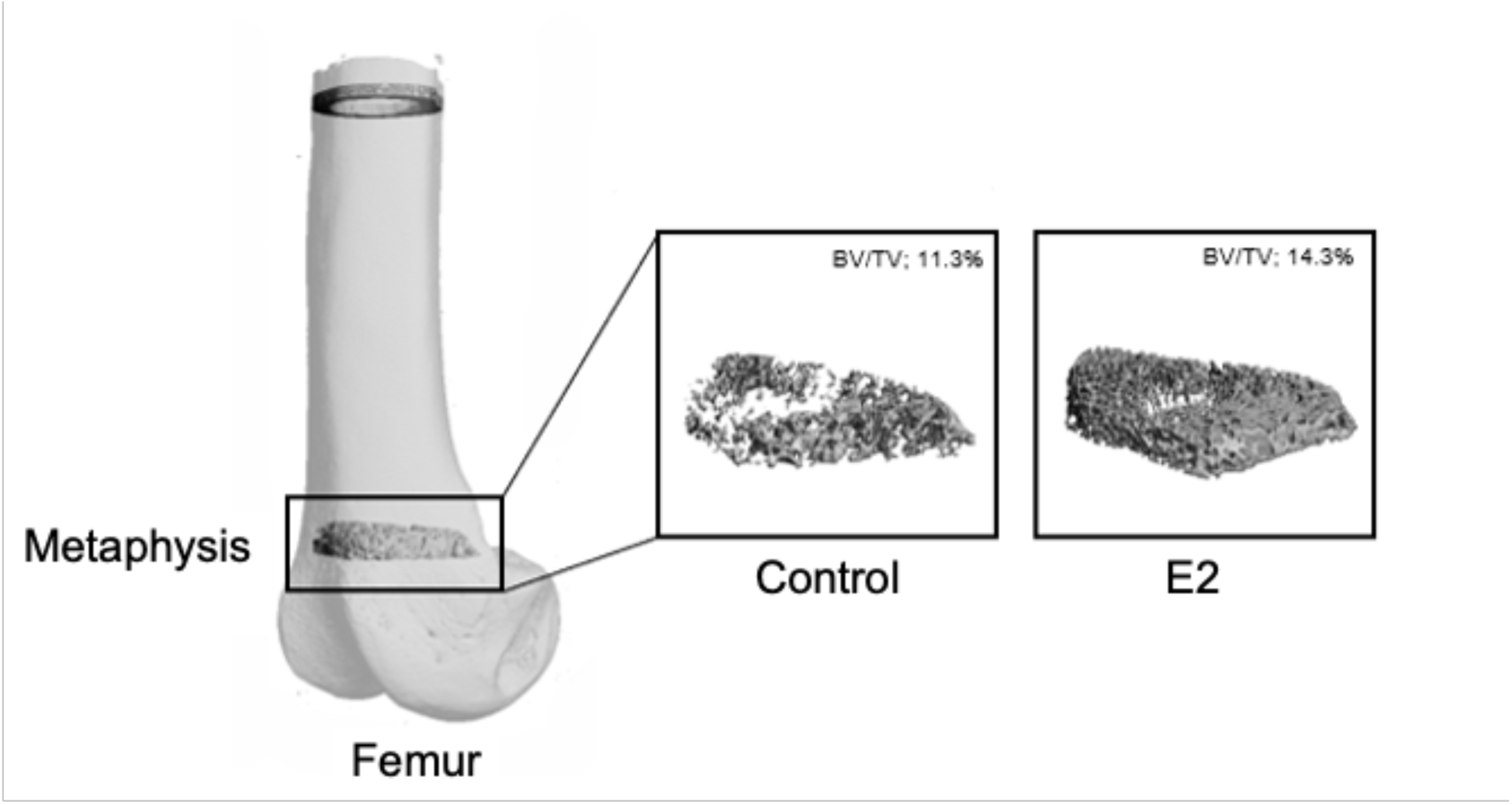
Representative μCT images of cancellous bone density (BV/TV) at the metaphyseal region of the femur in E2-replaced versus control animals.

**Table 2.**
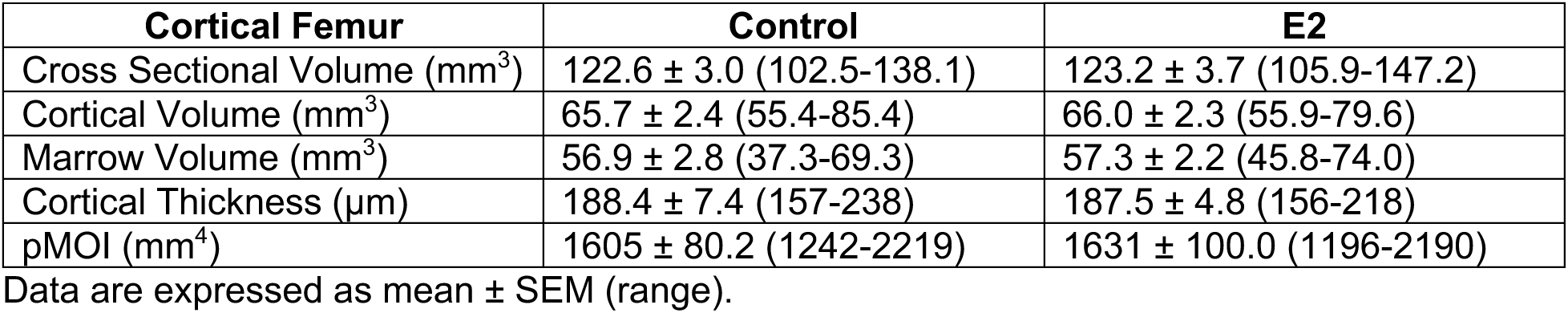
Structural bone parameters in cortical femur.

**Table 3.** Structural bone parameters in cancellous tibia.

| Cancellous Tibia | Control | E2 |
| --- | --- | --- |
| Bone Volume/Tissue Volume (%) | 11.1 ± 1.0 (7.6-16.7) | 13.3 ± 1.1 (8.1-22.9) |
| Connectivity Density (mm<sup>-3</sup>) | 3.7 ± 0.5 (1.4-6.5) | 5.2 ± 0.7 (2.2-11.1) |
| Trabecular Number (mm<sup>-1</sup>) | 1.3 ± 0.07 (1.0-1.7) | 1.5 ± 0.08 (1.0-2.1) |
| Trabecular Thickness (μm) | 144.2 ± 6.3 (122-198) | 143.7 ± 4.7 (124-173) |
| Trabecular Spacing (μm) | 781.4 ± 41.6 (600-1020) | 685.9 ± 37.2 (450-968) |

Data are expressed as mean ± SEM (range).

We also examined cancellous bone of the lumbar vertebrae. The effects of E2 replacement on cancellous bone in response to treatment were similar to those seen in the femur, with significantly higher connectivity density and trabecular number with lower trabecular spacing and no change in trabecular thickness. Unlike in the femur, however, bone volume/tissue volume in E2-replaced animals only showed a trend (p = 0.07) towards a higher ratio (**Figure 14 F-J**).

To complement the structural analyses described above, we assessed the circulating levels of several biomarkers of bone metabolism, namely osteocalcin, C-terminal telopeptide of type 1 collagen (CTX), total 25-OH vitamin D, and parathyroid hormone (PTH). Osteocalcin is secreted by osteoblasts and levels in blood reflect global bone formation while CTX is a marker of global bone resorption. Osteocalcin and CTX levels were significantly increased following OVX in control animals, an effect that was prevented by E2 replacement (**Figure 16A,B**). Vitamin D and PTH play important roles in mineral homeostasis and bone metabolism. However, no significant changes were seen in either over the course of the study (**Figure 16C,D**).

**Figure 16.**
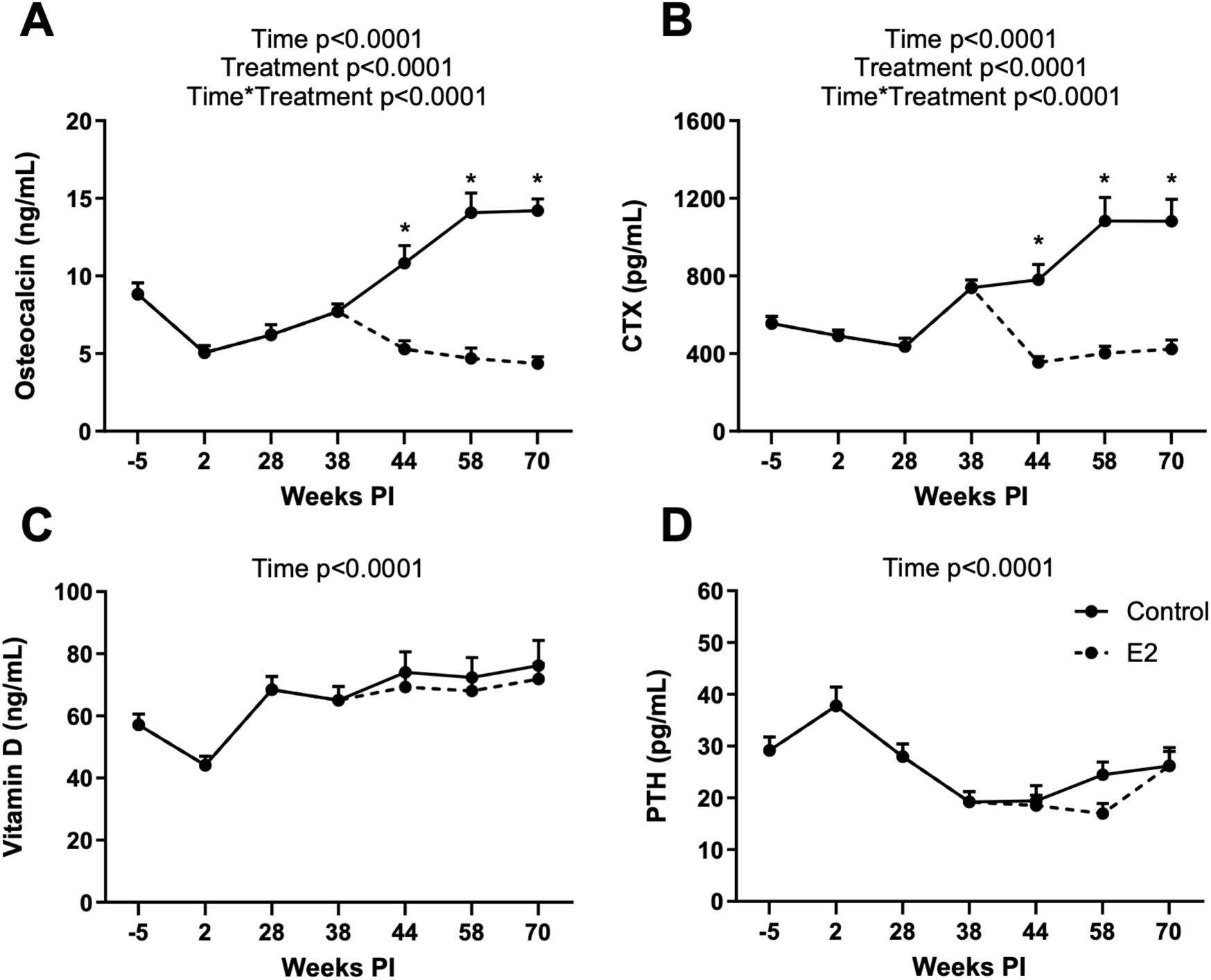
Effect of SIV infection, ART suppression, and E2 replacement on biomarkers of bone metabolism. Longitudinal changes in plasma (A) osteocalcin, (B) CTX, (C) total 25-OH vitamin D, and (D) PTH. All data are means ± SEM. Significance determined by mixed effects analysis with post-hoc pair-wise comparisons. *, p < 0.05.

Overall, these data support the conclusion that E2 exerts positive effect on bone health even in the context of SIV infection, ART treatment, and persistent inflammation.

## Discussion

Successful ART regimens have allowed many WLWH to survive through menopause and thus deal with the same consequences of hormonal (i.e., E2) depletion faced by women without HIV. The objective of this study was to assess the physiological changes resulting from ovarian hormone deficiency and the extent of their prevention by E2 replacement in the context of HIV infection and ART. This was accomplished by employing an NHP model of surgical menopause and E2 replacement. Our goals were to ascertain the effect of E2 deficiency and replacement on the viral reservoir in light of previous data implicating ERα in control of the latent reservoir (27,28) and to determine if previously described beneficial effects of E2 supplementation in women without HIV were preserved in the context of infection and ART and persistent inflammation that is not completely resolved by suppression of viremia.

With respect to the relationship between E2 and the HIV reservoir, several studies have suggested that E2 status may play a role in HIV infection per se (45–51) potentially through ERα inhibition of HIV transcription (24–26). A recent report suggests that E2 regulation of HIV transcription and, therefore, the HIV latent reservoir, may instead involve noncanonical ER signaling (52). Regardless of the specific molecular mechanism, we anticipated that maintenance of suppression of the cell-associated viral reservoir would be attenuated in E2 deficiency but maintained with E2 replacement. Instead, we observed no differences between the control and E2-replaced groups in either plasma viral loads or the cell-associated viral DNA levels in lymph nodes, colon, and WAT. These data suggest that E2 status does not exert a significant effect on the tissue reservoir in this in vivo model. Alternatively, it is possible that other mechanisms operative in vivo are sufficient to maintain viral suppression by ART despite a potential loss of E2 inhibition of viral transcription following menopause. We acknowledge that our determination of the tissue reservoir was based on the levels of total SIV DNA rather than measurement of inducible or replication-competent virus. However, we initiated ART during peak viremia at 2 weeks PI, and Long et al. (53) have previously shown that >80% of SIV genomes in PBMCs assessed >9 months PI in animals in which ART was initiated within 1 month PI are intact. Thus, we consider it likely that the majority of viral genomes in our cell-associated viral load assay are intact, such that a significant effect of E2 deficiency on the latent reservoir would be detectable in our assay.

Other parameters that did not exhibit changes based on E2 status in SIV-infected, ART-treated animals included SIV-specific humoral responses, peripheral and WAT immune cell profiles, systemic and islet-specific aspects of glucose homeostasis, lipid levels, and WAT morphology.

Cytokine profiling of plasma from a subset of E2-replaced animals revealed that, in addition to a significant increase in inflammatory cytokines following acute infection, a significant number of cytokines remained elevated following ART suppression of both the plasma and tissue viral reservoirs in both control and E2-replaced groups. We and others have previously reported a chronic inflammatory state following effective ART suppression of plasma viremia in male macaques (44, 54), and our findings, along with those of Fell et al. (55) and Chen et al. (56), extend the presence of post-ART chronic inflammation to female macaques as well. Overall, these data suggest that post-menopausal WLWH will experience a chronic inflammatory state independent of their E2 status.

A major observation was the positive effect of E2 replacement on several parameters of bone health and metabolism in the context of SIV infection and ART. While DXA scans are the gold standard for monitoring post-menopausal bone loss and subsequent osteoporosis, analyzing site-specific trabecular bone microstructure is far more detailed. Historically, this required ex vivo analysis of bone biopsies or, more recently, peripheral quantitative computed tomography (pQCT) (57). Examination of pre and post-menopausal transilial biopsies have shown decreased BV/TV and trabecular number with increased spacing (58) as well as the beneficial effects of E2 on these menopausal outcomes (59). Aged macaques have lower BV/TV values specifically in the hind limb (60) that are also seen in younger OVX animals (61, 62). E2-replaced females had increased trabecular number and connectivity density with decreased trabecular separation at two separate cancellous bone sites (femur and LV) compared to control. E2 females also had higher whole-body BMC and BMD as well as increased pelvis BMC when compared to control at 70 weeks PI. These positive effects on bone were evident in spite of the persistence of a chronic inflammatory state evidenced by a sustained increase in a significant number of inflammatory cytokines following effective suppression of viremia. An intriguing observation was the apparent progressive loss of pelvis BMC starting at the initiation of ART, suggesting that ART per se may exert an adverse effect on this parameter, as reported previously in WLWH (29, 63).

Postmenopausal bone loss is associated with elevated bone turnover and a negative bone remodeling balance (64). The increases in plasma osteocalcin and CTX levels following OVX seen in our NHP model are consistent with elevated bone turnover. Similar increases in biochemical markers of bone turnover were reported in OVX NHPs without SIV infection (65). The reduction in osteocalcin and CTX levels following E2 treatment is consistent with suppression of bone turnover. Higher whole-body BMD and BMC in the E2-treated animals are consistent with an improvement in bone turnover balance. Taken together, these findings suggest that SIV infection and ART have little influence on the skeletal response to E2.

The clinical implications of this study are heightened by the recent removal of the black-box warning from menopausal hormone therapy regimens, including E2-alone and E2-containing formulations (66). This step will likely increase the use of hormone therapy in general by peri and post-menopausal women (ideally within 10 years of menopause onset or before the age of 60), and our findings suggest that this would be appropriate for WLWH as well. E2 is typically delivered through oral or transdermal routes. The silastic implants employed in this study are more equivalent to the transdermal route (67), but transdermal and oral E2 replacement have been shown to result in similar beneficial effects on bone health (68). A notable strength of our study design is that the silastic implants we used resulted in serum E2 concentrations in the target range for the normal follicular phase.

We acknowledge that our study has some specific limitations. The first is our use of experimental (OVX-induced) menopause rather than natural menopause. The justification for this is that female macaques undergo natural menopause much later in their natural lifespan with fewer healthy years after ovarian failure (69, 70). In addition, NHP research facilities have relatively few healthy aged females, making the study of natural menopause impractical and poorly translatable. A second limitation is that we investigated the effects of E2 replacement alone rather than the effects of a combined E2-progestogen regimen, so that the effects of E2 in WLWH, particularly on bone health, who are also taking progestogens may differ from those we observed in the macaque model. An additional question is whether the effects we report in SIV-infected animals on a standard daily three-drug regimen of emtricitabine, dolutegravir, and tenofovir disoproxil fumarate would also be seen in WLWH who switch to newer long-acting ART formulations whose components may have novel modes of action, such as islatravir and lenacapavir (71, 72). Finally, sample size represents a limitation of our study, as we report multiple outcomes with statistically insignificant findings; however, the effect sizes we see do not support clinically meaningful differences in those parameters.

## Methods

### Sex as a biological variable

This study investigates the role of E2 replacement on viral dynamics and various aspects of metabolic heath in the context of post-menopausal E2 deficiency and thus required the use of cycling female macaques at study initiation.

### Study approval

All studies were approved by the Oregon National Primate Research Center Institutional Animal Care and Use Committee (IACUC) and were conducted in accordance with the US Public Health Service’s Policy on Humane Care and Use of Laboratory Animals.

### Animals

This study employed a total of 23 SIV-naïve adult female rhesus macaques, 4 of which (2 vehicle controls and 2 E2-replaced) were singly positive for the protective alleles *Mamu-B*17* or *Mamu-B*08* that are associated with spontaneous elite control of SIV (73). However, these animals did not exhibit any differences from the others with respect to viral dynamics. All animals were transferred from a regular chow diet to a Western-style diet (WSD) containing 36% calories from fat (Purina 5LOP, LabDiet) at least 8 weeks before infection and prior to baseline assessments at week −4 and maintained on the WSD for the remainder of the study. All animals were weight-stable at the time of baseline assessments. The rationale for the use of the WSD was the fact that a significant proportion of WLWH in the US consume a WSD and a large proportion are overweight or obese as a consequence of the global obesity epidemic (74–76). Animals were pair-housed in the same treatment group and maintained on a 07:00-19:00 light cycle with ad libitum access to water. Individual enrichment devices were provided in each cage, and animals were given weekly access to additional enrichment activities (radio, TV, etc.). All sedations for procedures were performed with ketamine HCl (5-15 mg/kg) (Covetrus, Dublin, OH) or a tiletamine HCl and zolazepam HCL mixture (3-5 mg/kg) (Telazol™; Zoetis Inc., Kalamazoo, MI) unless otherwise noted. For logistical and budgetary reasons, study animals were grouped into two cohorts studied consecutively. However, this schedule had the advantage of providing a randomized complete block design (77) recommended to improve precision of estimated effects. The first cohort consisted of 12 animals; however, one animal was excluded from the study due to detectable estrogen levels post-OVX. The second cohort consisted of 12 animals.

### Study design

The experimental plan is shown in **Figure 1A**. Baseline assessments were performed at 4 weeks before infection, with weekly blood draws commencing at infection (week 0). OVX was performed in all animals at week 35 PI and implants done at week 38 PI. All animals were necropsied at week 72 PI.

*WAT, lymph node, and colon biopsies, SIV infection, determination of plasma and cell-associated viral RNA levels, ENV-binding antibodies and SIV-specific T-cell responses, PBMC and WAT immune cell profiling, isolation of WAT immune cells and WAT histology, ivGTTs, determination of plasma leptin, adiponectin, HbA1c, and lipid levels, and necropsy.* All these procedures were performed as previously described (44). Antibodies used in PBMC and WAT immune cell profiling by flow cytometry are listed in **Table 4** and the gating strategy is shown in **Figure 17**.

**Figure 17.**
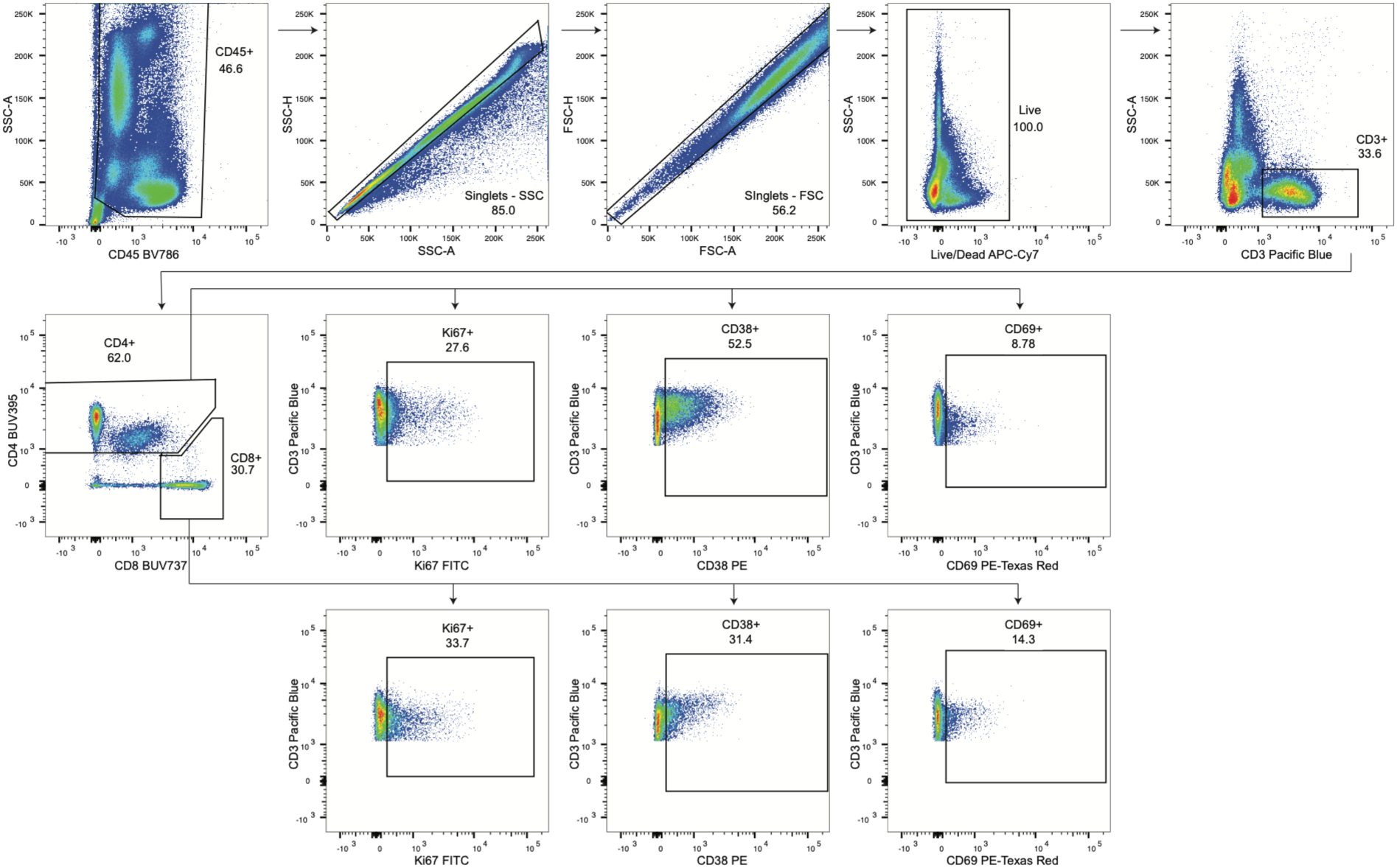
Gating strategy employed in immune cell profiling by flow cytometry. Top panels show steps in isolation of CD4+ and CD8+ T cells and bottom panels show steps in assessment of T cell activation based on expression of Ki67, CD38, and CD69.

**Table 4.**
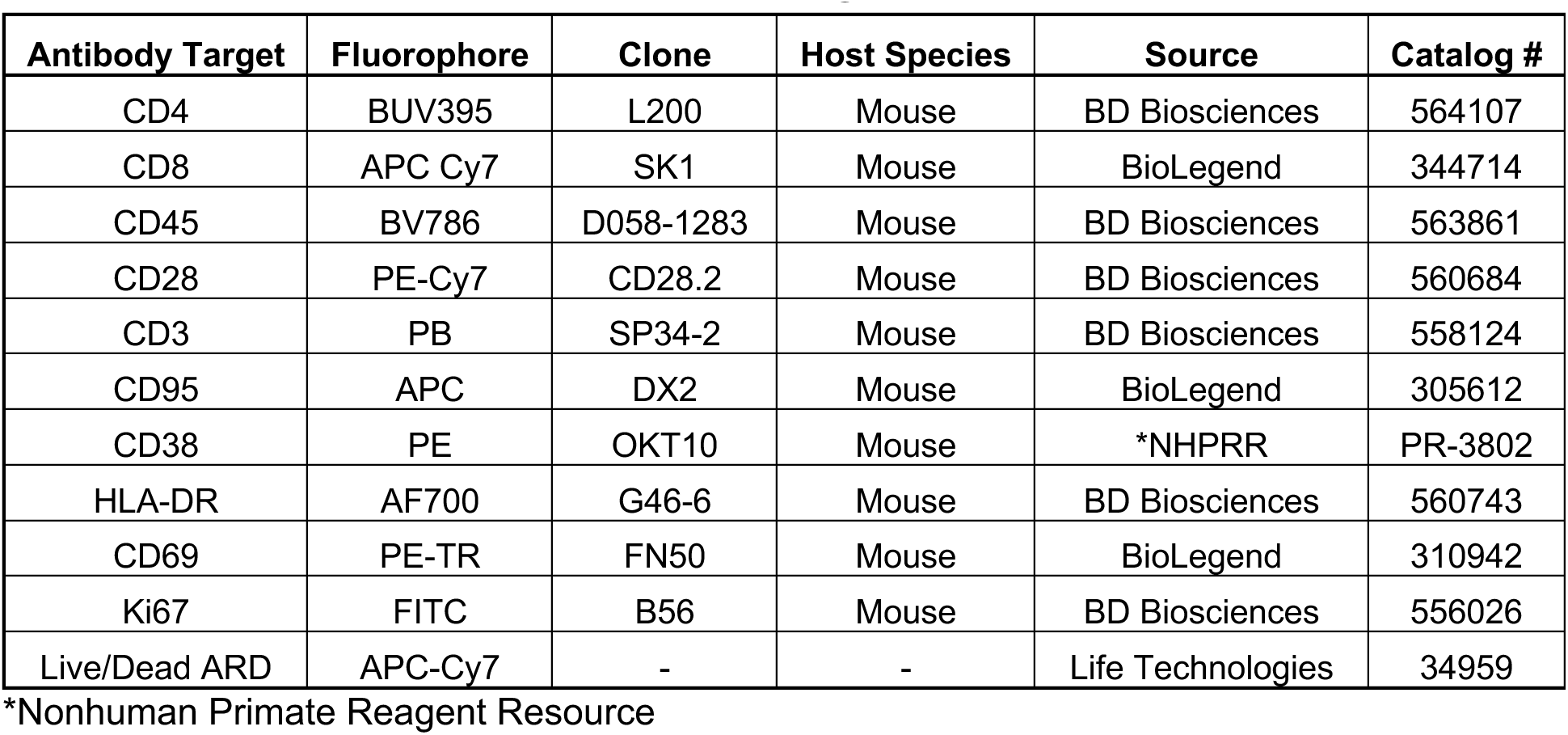
Antibodies used in immune cell profiling.

### ART regimen

At 2 weeks post-infection, an ART regimen consisting of dolutegravir, tenofovir disoproxil fumarate, and emtricitabine, co-formulated as a once-daily injection as described previously (78, 79), was initiated and continued for a subsequent ∼70 weeks.

### OVX and E2 replacement

After 4 months of ART, sufficient for acquisition of full suppression (**Figure 1B**), all animals were subjected to laparoscopic OVX at week 35 PI and implantation of silastic pumps to deliver E2 or cholesterol vehicle as described previously (31) at week 38 PI. Specifically, crystalline E2 (1,3,5,10-estratrien-3,17-β-diol; Steraloids, Wilton, NH) was administered via Silastic capsules (Dow Corning, Midland, MI) implanted subcutaneously in the periscapular region. Serum E2 was measured every week by ONPRC Endocrine Technologies Core in a Cobas e411 analyzer (Roche Diagnostics, Indianapolis, IN) with a sensitivity limit of the assay of 5 pg/mL. Dosing was adjusted to achieve target physiological levels of 50 to 125 pg/mL (**Figure 18**). If an individual’s concentration fell outside that range in two consecutive measurements, the capsule was replaced.

**Figure 18.**
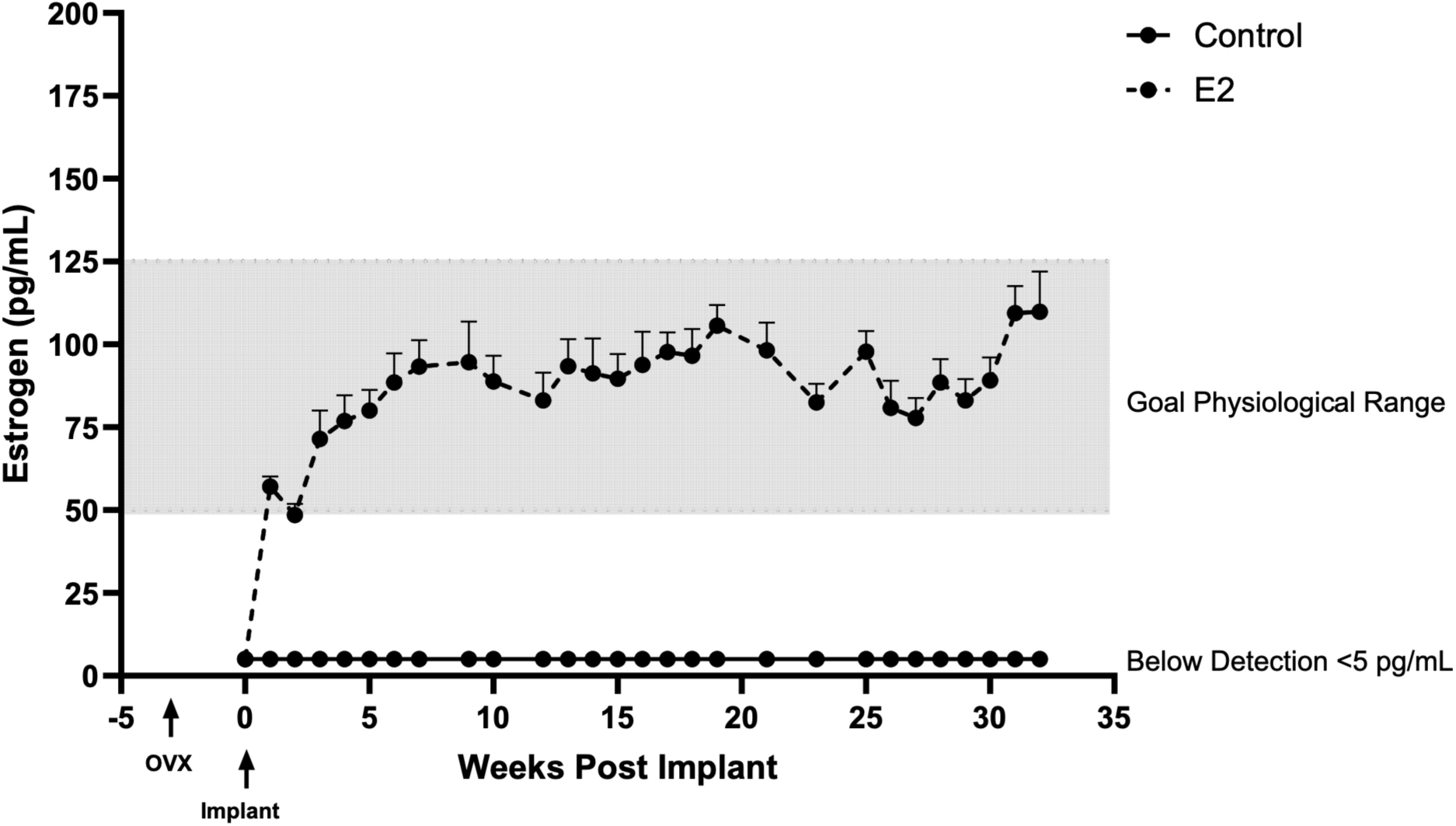
E2 levels in animals following E2 or control implants. Weekly plasma E2 levels in control and E2-replaced animals after OXV and implant placement. All data are means ± SEM.

### Body composition

Determination of total, lean, and fat mass as well as BMC and BMD were assessed using DXA scanning (Lunar iDXA, GE HealthCare, Chicago, IL). After an overnight fast, animals were sedated with 3-5 mg/kg Telazol and positioned in a prone position during a standard scan. These scans were typically paired with metabolic tests to decrease the number of sedations.

### Cytokine profiling

Longitudinal plasma samples (0, 2, 28, 38, and 58 weeks PI) from 6 E2-replaced animals were profiled at Alamar Biosciences, Inc. (Fremont, CA) using the NULISAseq^TM^ Inflammation panel that queries the circulating levels of 247 cytokines, 164 of which are detectable in macaque plasma (B. Burwitz, pers. comm).

### Islet isolation/perifusion

Islet isolation and perifusion assessment of glucose-stimulated insulin secretion was performed as previously described (44) with modifications. An 18-gauge plastic iv catheter was inserted into the pancreatic duct, and the pancreas was inflated with 50-80 mL of cold Hanks’ balanced salt solution with calcium and magnesium (HBSS; cat# 24020117, Gibco, Grand Island, NY) containing 10% heat-inactivated FBS, 1 μg/mL DNase I (Cat# 11284932001, Millipore Sigma, St. Louis, MO), and 0.6 mg/mL collagenase P (Cat# 11213865001, Millipore Sigma). After inflation, adipose and connective tissue was removed, and the pancreas was divided into 12 equal portions, with each portion placed in a 50-mL conical tube containing 11 mL of warm HBSS containing 0.06 mg/ml collagenase P and incubated in a 37°C water bath for 25 minutes with agitation every 10 minutes. After digestion, collagenase was pipetted off, and 10 mL of cold media (RPMI 1640, cat# 11875-085, Thermo Fisher Scientific, Waltham, MA), 10% HI-FBS, and 4 ug/mL DNase I was added and manually shaken for 45 seconds to disrupt remaining connective tissue. Undigested material was removed by straining through a 500-μm filter, brought to 200 mL with cold media, and centrifuged at 50 x g for 2 minutes at 4°C. The pellet was washed and centrifuged 3 times with cold media. Islet purification was performed using a discontinuous density gradient (Optiprep Density Gradient Medium, cat# D1556-250ML, Millipore Sigma) diluted with cold media. Digested material was resuspended in a 25% gradient solution, with 20%, 18%, and 11% gradient solutions layered sequentially above with a bulb pipet. The layered gradient was centrifuged with no brake at 500 x g for 5 minutes at 4°C. The islet layer was collected and washed in 50 mL cold media, and an islet subset was stained with dithizone (Cat# 43820, Millipore Sigma) in 0.5% DMSO in PBS to assess quantity and purity. Islets were plated on 5% HI-FBS-precoated 10-cm plates in 50 mL cold media warmed to 37°C and cultured overnight at 37°C/5%CO2.

After overnight recovery, islets were transferred into prepared columns and placed in a perifusion system (Model PERI-4.2, Biorep Technologies, Miami Lakes, FL) maintained at 37°C. Islets were pre-incubated in Krebs-Ringer bicarbonate HEPES buffer (KRBH) containing 4.0 mM glucose for one hour at a flow rate of 100 ul/min. After pre-incubation, islets underwent alternating 15-minute washes in 4.0 or 16.7 mM glucose-supplemented KRBH for a total of 45 min, with collections of flow-through every three minutes. All perifusion assays were done in triplicate. Collections were stored at 4°C until being assayed for Insulin by ELISA as described above.

### Islet oxygen consumption

After overnight recovery, islet oxygen consumption was assessed using the Agilent Seahorse XF Mito Stress Test and XFe24 Analyzer (Agilent Technologies, Santa Clara, CA). 20-30 islets were selected per well, with 5 replicate wells, containing Seahorse XF RPMI medium (Cat# 103576, Agilent Technologies) supplemented with 4mM glucose (Agilent cat# 103577-100), 2 mM glutamine (Cat# 103579-100, Agilent Technologies), and 1% HI-FBS. The four ports of the XFe24 Islet Flux Plate were loaded with 200 mM glucose, 50 μM oligomycin A, 10 μM carbonyl cyanide ρ-trifluoromethoxyphenylhydrazone (FCCP), and 30 μM rotenone/antimycin A (Cat# 103015-100, Agilent Technologies), diluted in XF RPMI medium. The Wave assay protocol was programmed with a 24-minute basal period, a 48-minute high-glucose period, a 48-minute oligomycin period, a 32-minute FCCP period, and a 48-minute rotenone/antimycin period, with oxygen and pH measurements taken every 8 minutes.

### μCT analyses

μCT was used for nondestructive 3-dimensional evaluation of bone architecture in femur, tibia, and lumbar vertebra. Distal femora, proximal tibiae, and 3rd lumbar vertebrae were scanned in 70% ethanol at a voxel size of 36×36×36 μm (55 kV p, 145 μA, and 200 μs, 500 projections/rotation) on a Scanco μCT40 scanner (Scanco Medical AG, Bassersdorf, Switzerland). Filtering parameters sigma and support were set to 0.8 and 1, respectively. Bone segmentation (differentiation of bone from non-bone) was determined empirically and set at a threshold of 175 (scale, 0-1000). Cortical bone was evaluated in thirty consecutive slices (1.1 mm) starting at the proximal end of the distal third of the femur. Cancellous bone was evaluated in the distal femur metaphysis, proximal tibia metaphysis, and vertebral body. The region of interest in the femoral metaphysis consisted of 60 slices (2.2 mm) proximal to the superior part of the patellar surface. The region of interest in the tibial metaphysis consisted of 60 slices (2.2 mm) distal to the lowest extent of the medial condyle. The entire cancellous envelope (536 ± 7 slices; 19.3 ± 0.3 mm) was evaluated in the vertebral body. Irregular manual contouring a few pixels interior to the endocortical surface was used to delineate cancellous bone from the endocortical surface. Cortical measurements included (1) cross-sectional volume (cortical and marrow volume, mm^3^), (2) cortical volume (mm^3^), (3) marrow volume (mm^3^), (4) cortical thickness (μm), and (5) polar moment of inertia (a surrogate measure of bone strength in torsion, mm^4^). Direct cancellous bone measurements included cancellous bone volume fraction (bone volume/tissue volume, BV/TV, %), connectivity density (mm^-3^), trabecular number (mm^-1^), trabecular thickness (μm), and trabecular spacing (μm).

### Circulating biomarkers of bone metabolism

Plasma levels of osteocalcin, CTX, total 25-OH vitamin D, and PTH were assayed by the ONPRC Endocrine Technologies Core using a Roche Cobas e411 analyzer (Roche Diagnostics, Indianapolis, IN).

### Statistics

Statistical analyses were performed using GraphPad Prism (GraphPad Software, Boston, MA). AUC was calculated starting at T=0 using the trapezoidal rule. Longitudinal data for parameters that had baseline values were analyzed for effects of time (Time in figures), intrinsic differences between the groups (Treatment in figures), and differences in the longitudinal response between the groups (Time*Treatment in figures) using mixed-effect analysis with post-hoc multiple comparisons test. Correlations were determined using Pearson’s correlation coefficient. A p value <0.05 was considered significant.

## Data availability

All supporting data will be provided by the corresponding author upon request.

## Author contributions

JBS, PK, CTR, and KAS designed the study, GMW, DT, MK, SRL, HB, CZ, MS, CM, HF, CP, AM, HM, VM, and OV conducted experiments and acquired data. GMW, UTI, RTT, OV, PCT, JTJ, JBS, PK, CTR, and KAS interpreted the results. GMW, CTR, and KAS wrote the paper. All authors reviewed and edited the manuscript.

## Acknowledgements

We acknowledge the assistance of the ONPRC flow cytometry, molecular virology, endocrine technologies and integrated pathology cores. This study was supported by NIH grants R01 AG07114 to JBS and PK, K01 AI183927 to KAS, P51 OD011092 for operation of the ONPRC, and S10 OD25002 for support of the Aperio AT2 instrument employed for analysis of WAT histology.

